# Biclique extension as an effective approach to predict novel interaction partners in metabolic compound-protein interaction networks

**DOI:** 10.1101/2021.09.23.461460

**Authors:** Sandra Thieme, Dirk Walther

## Abstract

**Motivation:** Metabolic networks are complex systems of connected chemical reactions with physical interactions between metabolites and proteins playing a critical role for both metabolic conversion and regulation. In this study, we aimed to predict previously unknown compound-protein interactions (CPI) by transferring the concept of biclique extension, which was developed in the context of drug-target interaction prediction and that is based on the rationale that interactions that readily extend an existing biclique are real, to metabolic CPI networks.

**Results:** We developed and tested a workflow to predict CPIs based on the concept of extending existing bicliques and applied it to *E. coli* and human using their respective known CPI network as input. Depending on the chosen biclique size, for the *E. coli* network we reached a sensitivity of 39% with an associated precision of 59%. For the larger human CPI network, a sensitivity of 78% with a false-positive rate of less than 5% and an associated precision of 75% was obtained. At more stringent settings, a precision as high as 95% was attainable at the expense of a lowered recall. Prediction performance significantly exceeded that obtained using randomized networks as input. Predicted novel interactions were tested for biomolecular function involvement, with TCA-cycle and ribosomal processes found associated with particularly pronounced statistical significance. As we demonstrate, our approach holds great potential to increase efficiency of experimental testing of CPIs and can readily be transferred to other species of interest.

**Availability and implementation:** The R code and datasets are available at https://github.com/SandraThieme/BiPredict.

## 1. INTRODUCTION

Identifying novel compound-protein interactions (CPIs) is a central research objective of molecular biology as it can be considered critical for understanding biological systems at the molecular level. Recently published studies reported large-scale experimental screens for the identification of novel interactions between compounds and proteins (Piazza et al. 2018; Diether et al. 2019). In these studies, a large number of potential interactions was experimentally tested with only a relatively small fraction of candidate interactions being validated (around 5% for Piazza et al. 2018). Thus, augmenting experimental approaches with bioinformatic workflows may help narrow down the set of candidates for experimental testing, and, thus, increase the rate of successfully validated interactions, while also saving time and resources. This study aims to test the utility of computational approaches that employ the so-called biclique extension method to serve this goal.

Compound-protein interaction networks can be represented as a bipartite graph, in which nodes represent compounds and proteins as the dual entities or groups, and edges represent the interactions between them, but not between compounds or proteins themselves. In a bipartite network, a subset of nodes with the maximum number of possible connecting edges between them actually established is defined as a biclique. Thus, a biclique represents the densest possible connection between a subset of nodes in such a network. We aimed to use bicliques to identify such very closely related sets of compounds and proteins in known CPI networks, and to search for interaction candidates in the directly connected neighborhood of these bicliques. The concept is based on the logic that interactions between compounds and proteins are likely true, if, by postulating them, an existing biclique is expanded (see Figure 1 for a schematic illustration and further explanation of the underlying logic). Biclique-based approaches have been used in drug-target interaction (DTI) networks for the prediction of novel pharmaceutically relevant chemicals or unknown target proteins (Daminelli et al. 2012), and for the prediction of protein-protein interactions (Schweiger, Linial, and Linial 2011). Other recently published methods include neural networks based on chemical properties and structure information (Tsubaki, Tomii, and Sese 2019; Eslami Manoochehri and Nourani 2020) as well as random forests based on GO-terms and KEGG pathway enrichment in combination with chemical substructure information (Chen et al. 2016). Also bipartite local models (BLM) have been widely used for DTI prediction, for example, based on chemical and protein sequence similarity (Daminelli et al. 2015; Bleakley and Yamanishi 2009) or based on expression data in combination with localization information of enzymes and phylogenetic profiles (Bleakley, Biau, and Vert 2007). BLMs apply network-theoretic approaches for the prediction of interactions that employ local topological information (Cannistraci, Alanis-Lobato, and Ravasi 2013) to bipartite networks. For recent reviews on DTI predictions, see (Lotfi Shahreza et al. 2018) and (Z. Wu et al. 2018).

**Figure 1.**
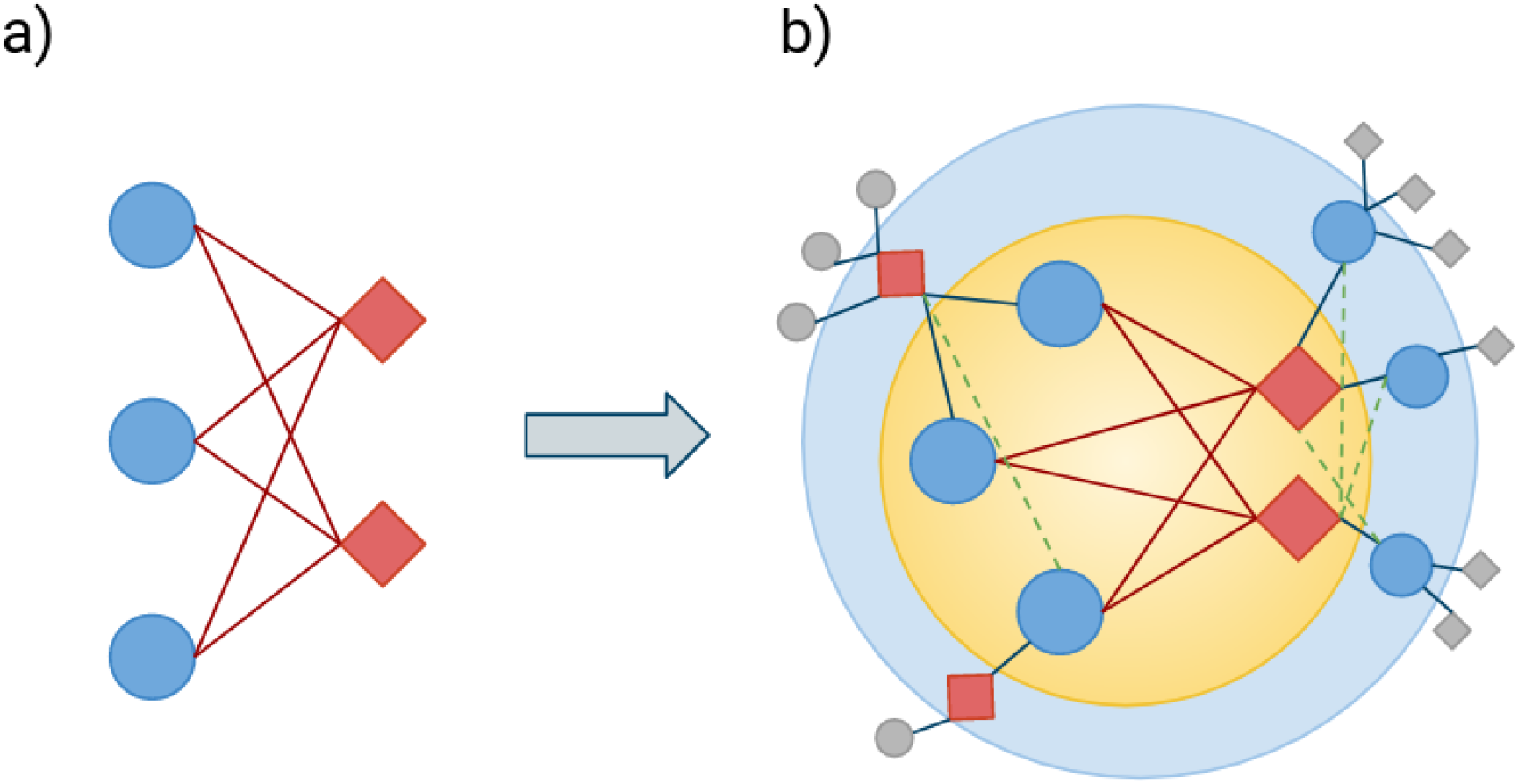
Biclique definition and extension. Bicliques consist of two types of nodes and edges connecting each node of different types (a). Here, blue circles represent compounds, c, and red squares represent proteins, p, while edges represent interactions. The biclique expansion starts with an existing biclique (b, yellow inner circle), here consisting of three compounds (c = 3) and two proteins (p = 2). Next, all compounds and proteins that are directly connected to any member of the biclique are identified. They represent interaction candidates (b, lightblue outer circle). Other compounds and proteins of the network which are not directly connected to any member of an existing biclique are not considered (b, grey squares and circles). Finally, interactions are predicted if interaction candidates lack only one edge to be a member of an existing biclique (b, green dashed lines).

We aimed to transfer the biclique extension approach from the field of DTI networks to the identification of as of yet unidentified CPIs in naturally occurring metabolic and cellular networks. The increasing number of experimentally verified interactions in public databases allows us now to search for new interactions solely on the basis of known interactions, without adding any other sources of information to our network. By focusing only on the network structure in our approach, we aimed to reduce the need for additional data, but also to limit the risks of overfitting caused by the high dimensional feature space, applied typically in previous approaches.

A crucial element for assessing the accuracy of CPI predictions is the availability of known negative interactions, which refers to CPIs that are confirmed to have no interaction under natural conditions (Liu et al. 2015). The validation of newly predicted interactions benefits from having both true-positive and true-negative interactions determined in wet-lab experiments, which are not yet part of the input interaction network the predictions are based on. In contrast to most published CPI prediction methods, we did not only use randomly generated negative samples for the verification of our predicted interactions, as it is difficult to assess for such random data how many positive interactions are actually contained in such random connections. Instead, we used negative interactions as reported experimentally.

We analyzed an *E. coli* CPI network to test the prediction performance on recently published experimental data and also applied the biclique extension method to a human CPI network for which a computed validation dataset based on a huge number of biological features was available. In addition to our biclique based predictions, we studied the network properties, which proved relevant for the biclique predictions.

## 2. METHODS

### 2.1 Overview

We established the following biclique extension workflow, summarized in Figure 2. First, a reference network of validated interactions was constructed. Here, we computed a compound-protein-interaction (CPI) network based on data from the STITCH database for *E. coli* and human. Next, we used annotation information from the KEGG database to remove all interactions of drugs from our network to obtain a naturally occurring metabolic/ cellular network. We predicted novel interactions based on interaction candidates, which were connected to existing bicliques in the reference network (Figure 1). For validation of our predictions, two additional datasets were needed. First, a true-positive dataset of true interactions, which are not part of the reference network and, secondly, a negative dataset for which no interactions could be shown to serve as a true-negative set. Here, we used datasets generated based on experimental data for the *E. coli* network (Piazza et al. 2018) and computationally predicted data for the human network (Liu et al. 2015) to evaluate our results.

**Figure 2.**
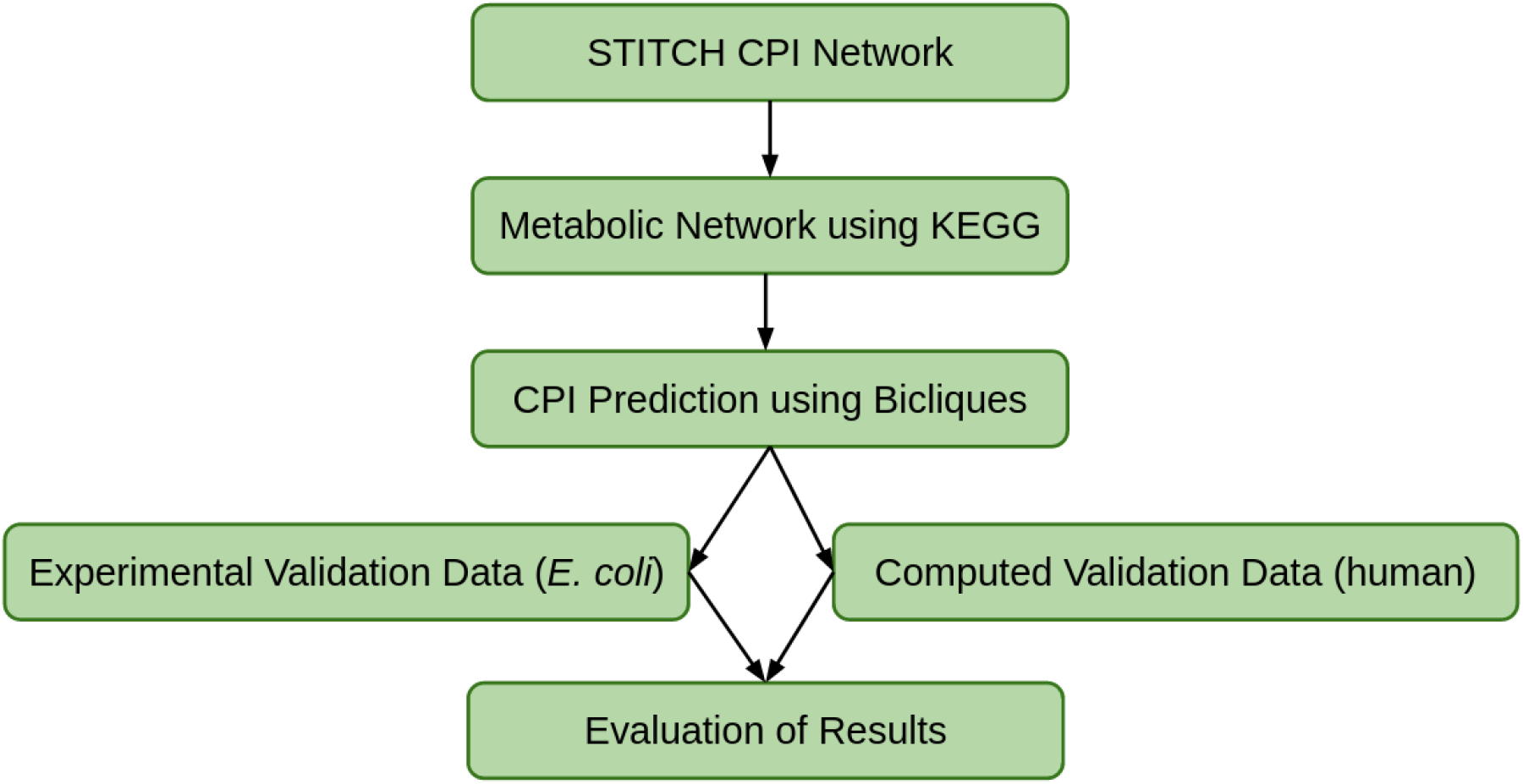
Biclique prediction workflow. We used a STITCH CPI network as input and removed all non-naturally occurring interactions information from KEGG to obtain a naturally occurring metabolic network. We predicted interaction candidates using bicliques and evaluated our predictions using different validation datasets derived from sampling, experimental and computational studies.

### 2.2 The Reference Network

Information about known and predicted interactions between compounds and proteins of *Escherichia coli K12 MG1655* and human was downloaded from the STITCH database (http://stitch.embl.de, version 5.0; Kuhn et al. 2008; Szklarczyk et al. 2016), including links to other databases, names, and SMILES strings of compounds. Protein sequences and links to other protein databases were downloaded from the STRING database (Szklarczyk et al. 2019), UniProt (The UniProt Consortium 2017), and from LINKDB (Fujibuchi et al. 1998). STITCH database entries of compounds were rendered non-redundant by merging identifiers capturing isoform and salt variants. STITCH also provides a confidence score for every reported interaction, ranging between zero and one and with larger values corresponding to higher confidence (qualitative intervals: low scores of 0.0-0.4, medium 0.4-0.7, high: 0.7-0.9, and very high confidence: 0.9-1).

Out of the total of 1,821,709 reported interactions for *E. coli*, 242,125 interactions were assigned a confidence of ‘medium’ or better. To infer the CPI network, only experimentally verified interactions were included, representing edges in our network (176,100 STITCH interactions). Both, direct as well as transferred confidence scores, i.e. scores assigned by homology from other species, were taken into account. We applied a ‘medium’ experimental confidence score of 0.4 on the STITCH network as a lower threshold, which included 99,487 out of 1,821,709 reported interactions. To test for robustness of our predictions, networks based on confidence score thresholds ranging between 0.4 and 0.6 were tested as well. Thus, we tested our predictions on three networks ranging in size from 37,655 (score>0.6) interactions to up to 99,487 (score>0.4) interactions.

Out of the total of 15,473,939 reported interactions in human, 1,545,933 interactions were assigned a STITCH confidence score of ‘medium’ or better. 8,842,952 STITCH interactions carry experimental support. We applied a ‘medium’ experimental (considering direct and transferred) confidence score of 0.4 as a lower threshold on the human network, which resulted in 1,026,207 interactions. To test the robustness of our predictions, a network based on a confidence score of 0.5, including 641,457 interactions was tested as well.

### 2.3 Network Construction

The bipartite CPI network was computed using the R package *igraph* (Csardi and Nepusz 2006; R Core Team 2016). Nodes of the network represent compounds and proteins. Edges were inserted connecting compounds and proteins, for which known interactions based on the filter criteria described above were reported.

### 2.4 Data Cleanup

To exclude unspecific interactions, small compounds such as ions were removed from the network. Correspondingly, all interactions with compounds of less than five heavy atoms, as determined from their SMILES strings, were removed from the initial dataset, as done similarly by Daminelli et al. 2012 (Daminelli et al. 2012). Also, the STITCH CPI network contains many interactions of compounds, which are not naturally occurring in the metabolic or cellular network of the corresponding organism, such as antibiotics. We used additional annotation information from KEGG (Kanehisa et al. 2017) to confine our sets of compounds to metabolites and naturally occurring cellular compounds (‘C’ number KEGG compounds). In addition, compounds marked as ‘antibiotics’, and (in *E. coli*) ‘hormones and transmitters’ or ‘steroids’ in KEGG were removed from the network. Also compounds with a drug ID in KEGG (‘D’ number) and without an additional ‘C’ number in KEGG were also removed from the network. Compounds assigned a ‘D’ number, but also a ‘C’ number were retained.

Starting with the STITCH *E. coli* network of 69,109 nonredundant interactions, 6,894 interactions were retained after data cleanup. The same data cleanup steps were applied to the human network resulting in a network of 44,322 interactions.

In addition, prior to the biclique calculations, we excluded all interactions with compounds and proteins occurring only once (i.e. one reported interaction only) from the network, because such interactions were not relevant for the biclique calculations, which we required to consist of at least two compounds and proteins (i.e. each biclique member must have at least two interactions). This reduced the E. coli network to 6,353 interactions and the human network to a size of 42,158 interactions.

### 2.5 Validation Data

The prediction performance of the biclique extension method was tested on two reference CPI networks from two species: *E. coli* and human. For both species, true-positive interactions were taken as random samples from high-confidence STITCH interactions.

True-negative interaction compound-protein-pairs were taken from two different resources. For *E. coli* we used interactions which were neither reported in STITCH nor detected interacting in a recent experimental assay in which the interactions were tested (Piazza et al. 2018; Diether et al. 2019).

For human, a true-negative set was taken from a study that aimed to computationally assemble a high-confidence negative CPI-set (Liu et al. 2015).

In both cases, only those validation-set-CPIs were considered, for which the corresponding compounds and proteins were found present in the STITCH reference network. The specifics of the validation sets are outlined below.

#### 2.5.1 Validation procedure and data, *E. coli*

As a positive validation dataset, in each of the ten performed prediction runs, ten percent of the true interactions as reported by the STITCH reference network were randomly chosen and considered predictable true-positives (hereafter referred to as *positives*). All *positives* were removed from the reference CPI network prior to the biclique detection and subsequent prediction. They were used to calculate the true positive rate (TPR) of predicted interactions.

A validation set of negative interactions was compiled from experimental data. Piazza *et al*. experimentally tested 34,186 interactions between 20 central compounds (e.g., ATP, ADP, NADP; Supplementary Table 1) and 2,525 proteins of interest and reported 1,719 interactions. A negative validation dataset for our study was created based on the 32,467 interactions, which were tested by Piazza *et al*., and reported to not interact in their experiments. A subset of 26,724 interactions consisted of compounds and proteins that were also part of the STITCH network and not known to interact (hereafter referred to as *Piazza.negatives*). This dataset was expanded by a second recently published interaction dataset that also reported experimentally tested interactions in *E. coli* (Diether et al. 2019). From the experimental data from Diether *et al*., an additional set of 1,354 tested, but reported as non-interacting compound-protein pairs comprising 55 compounds and 29 proteins was included as true-negatives in our study. However, 584 of these supposedly negative interactions were included in the STITCH database of known and predicted interactions. These interactions were removed from the validation dataset of negatives, which finally comprised 863 non-interacting compound-protein-pairs (hereafter referred to as *Diether.negatives*). By adding this dataset, the number of compounds included in the validation data (*negatives*) was increased from 20 to 57. As 172 interactions of *Diether.negatives* were already contained in the *Piazza.negatives*, the combined *negatives* list finally included 27,415 unique experimentally verified non-interactions between 57 chemicals and 2,474 proteins.

Depending on the chosen confidence-score of the reference STITCH network, about 10,720 of these negative interactions (*negatives*) were available for prediction; i.e. both the compounds and proteins were present in STITCH. They were used to calculate the false positive rate (FPR) of predicted interactions.

Only predictable interactions were taken into account to calculate true-positive (TP) and false-positive (FP) interactions and rates, i.e. only interactions between compounds and proteins, which were both included in the reference network, and, thus, could be predicted by biclique extension.

#### 2.5.2 Validation procedure and data, Human

A positive validation dataset was created by randomly choosing 5% of interactions from the input network as done in *E.coli*. Due to the larger network (44,322 interactions), the random set constituted 5% of true interactions in human, not 10% as done in *E.coli* (6,894 interactions).

A dataset of negative interactions was taken from Liu et al. (Liu et al. 2015). This study published highly reliable negative interaction sets combining various chemical, structural, and interaction information. From the provided negatives dataset, 39,758 out of 40,381 compounds and 1,974 out of 2,027 proteins could be successfully mapped to STITCH IDs, yielding a total of 369,276 negative interactions. Of these, 15,865 were predictable negative interactions (*negatives*) with chosen confidence-score of the reference STITCH network of 0.4. All interactions between compounds and proteins that were also included in the input network were considered to be predictable interactions.

### 2.6 Biclique Calculation and Extension

All maximum bicliques of compounds and proteins in the CPI network were calculated using the R package *biclique* (Lu, Phillips, and Langston 2020; Yun Zhang et al. 2014). Using maximum-size bicliques makes sure that no bicliques of a certain size that are fully contained in larger bicliques are considered separately. However, overlapping bicliques are possible, and therefore, edges, representing interactions between two nodes, can be members of multiple bicliques.

A minimum number of two and up to nine nodes on either side of the biclique was tested, representing the minimum number of compounds and proteins of each biclique. Thus, the smallest considered bicliques consisted of two proteins and two compounds.

Candidates for novel interactions between compounds and proteins in the network were searched in the directly connected neighborhood of existing biclique-member nodes, i.e. proteins and compounds, which are connected to at least one node of the biclique. Only compounds and proteins, which become part of an existing biclique by insertion of exactly one connecting edge, were considered as novel interaction candidates (see Figure 1 for a schematic illustration). In addition, an insertion of two edges was tested for bicliques with more than four nodes on the corresponding side.

### 2.7 Molecular similarity measures

The similarity of compounds was estimated based on the Tanimoto index with structural features derived from the SMILES string and using the R package *RxnSim* (Giri et al. 2015). The similarity of proteins was assessed based on pairwise protein-based BLAST alignment scores. *E. coli* protein sequences were downloaded from UniProt, ID: UP000000625, strain K12. Given the set of proteins included in the *E. coli* dataset, an all-against-all blastp search was performed with the E-value threshold set to 10 to allow for weak alignments to be reported and considering one (the best) HSP per pair only. Otherwise, default blastp settings were used. Cohen’s-d effect sizes were calculated using the R package effsize (Torchiano 2016).

### 2.8 Randomization

To investigate the importance of the underlying reference network of experimentally verified interactions for our predictions, we created a randomized network altering the reference interaction network. To generate a random bipartite network, the R package *BiRewire* (Gobbi et al. 2020) was used. *BiRewire* uses the edge switch algorithm to preserve node degrees of the input network. This randomized network was also used for biclique calculation and extension.

### 2.9 Prediction performance metrics

Prediction results were assessed with regard to true-positive rate (TPR, or sensitivity or recall), false-positive rate (FPR), and F1-measure, and precision as commonly defined.

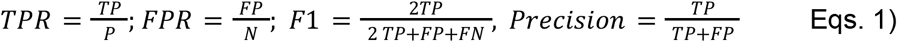

where P is the number of positives, N - the number of true negatives (no interaction), TP - true positive predictions, FP - false-positive predictions, and FN - false-negative predictions, assessed based on the positive and negative validation datasets described above.

### 2.10 KEGG Enrichment Analysis

To inspect our prediction results in terms of biological function and their biochemical processes, we performed a KEGG enrichment analysis on the *E. coli* network using the R package clusterProfiler (Yu et al. 2012; T. Wu et al. 2021).

### 2.11 Code availability

The developed R-script along with relevant data used in this study is available at https://github.com/SandraThieme/BiPredict

## 3. RESULTS

### 3.1 Reference Network Properties and Structure

To apply biclique extension to predict novel compound-protein interactions, we first compiled an *E. coli* reference interaction network based on data from the STITCH database. The *E. coli* network included 6,894 interactions between 177 compounds and 1,906 proteins, and after removing degree-one interactions (as they are not relevant for our method), 6,353 interactions between 160 compounded and 1,381 proteins (Table 1). The maximum node degree of compounds was 1,339 (glycerol), the mean degree was 38.9 and the median degree was 13. The maximum degree of proteins was 19 (gudD,(D)-glucarate dehydratase 1), the mean degree was 3.6 and the median degree was 3.

**Table 1.**
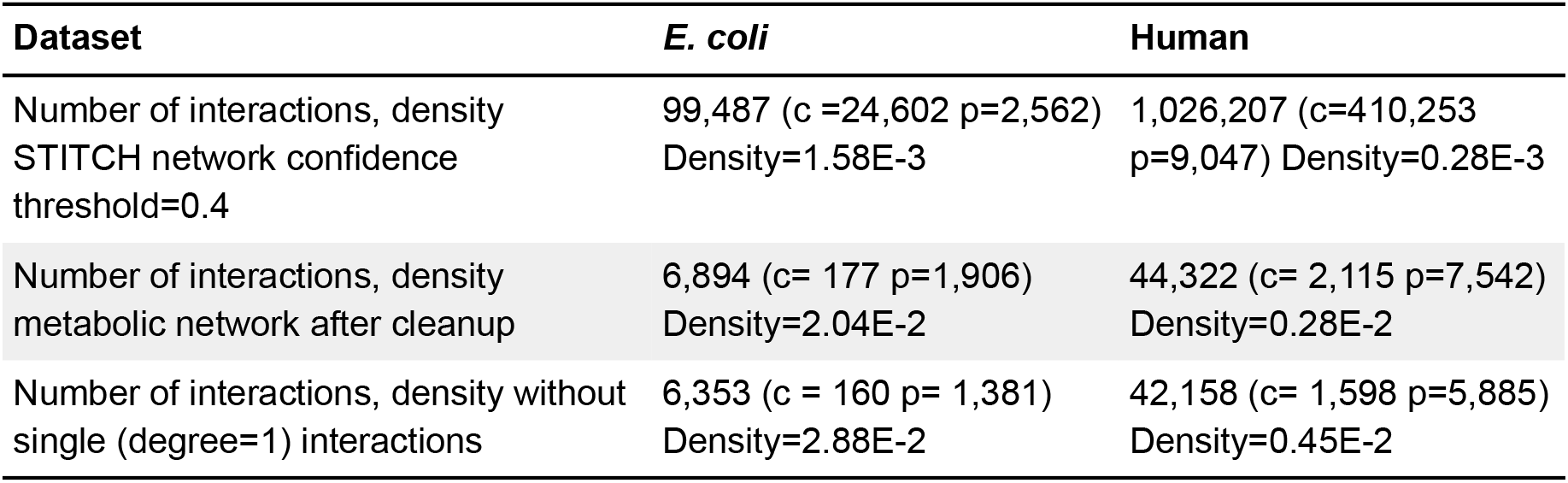
Number of interactions (*N_i_*), compounds (c) and proteins (p), and network density (=*N_i_*/(c*p) in the E. coli and human network after different filtering steps. Note, as node degree=1 interactions were not considered in the biclique computations (as they cannot contribute to our predictions), we list them separately.

| Dataset | *E. coli* | Human |
| --- | --- | --- |
| Number of interactions, density<br>STITCH network confidence<br>threshold=0.4 | 99,487 ($c=24,602$ $p=2,562$)<br>Density=1.58E-3 | 1,026,207 ($c=410,253$<br>$p=9,047$) Density=0.28E-3 |
| Number of interactions, density<br>metabolic network after cleanup | 6,894 ($c=177$ $p=1,906$)<br>Density=2.04E-2 | 44,322 ($c=2,115$ $p=7,542$)<br>Density=0.28E-2 |
| Number of interactions, density without<br>single (degree=1) interactions | 6,353 ($c=160$ $p=1,381$)<br>Density=2.88E-2 | 42,158 ($c=1,598$ $p=5,885$)<br>Density=0.45E-2 |

The human network consisted of 44,322/ 42,158 interactions including 2,115/ 1,598 compounds and 7,542/ 5,885 proteins, with and without degree-one interactions, respectively (Table 1). The maximum node degree of compounds was 2,959 (selenomethionine), the mean degree was 20.9 and the median was 3. The maximum degree of proteins was 100 (5-hydroxytryptamine (serotonin) receptor 2A), the mean degree was 5.9 and the median 3. Judging by their network density, the *E. coli* network had a much higher network density than the human network (7/6-fold difference, Table 1 with/without degree=1 interactions).

Despite their different sizes, with many more interactions reported for human than for *E. coli*, both networks showed similar degree distributions when recorded for compounds and proteins, respectively (Figure 3). For a compound-centric view, both networks follow a power-law (linear relationship in log-log scale, Figure 3, left panel), with the human data shifted to higher counts due to its larger network size. Power-law degree distributions have been found to be a characteristic of biological networks (Lima-Mendez and Helden 2009). By contrast, a power law was less obvious for a protein-centric degree distribution (Figure 3, right panel) with counts dropping faster than expected from a power law alone.

**Figure 3.**
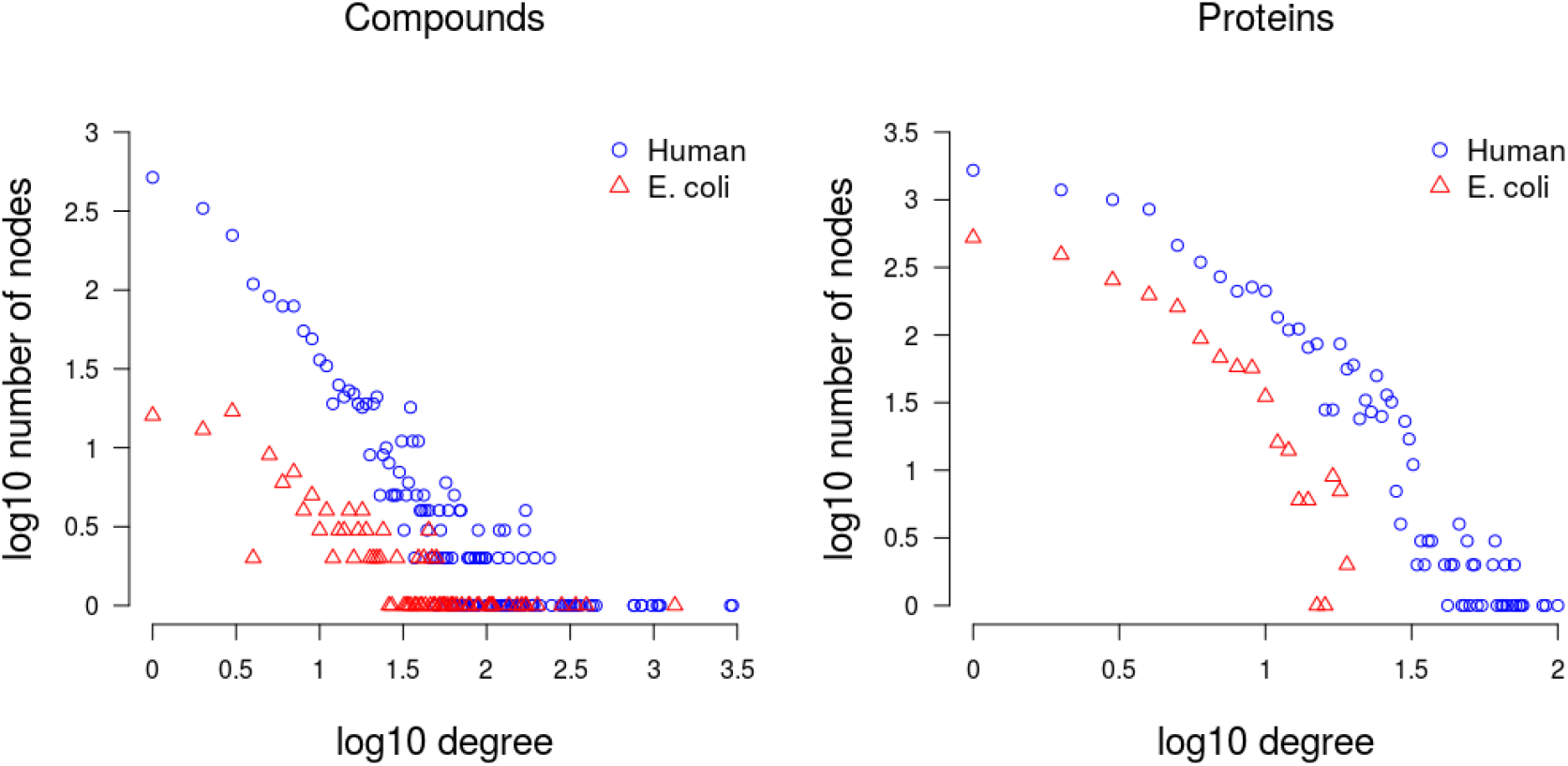
Degree distribution of compounds, (left panel), and proteins, (right) in the E. coli and Human STITCH reference network. The x-axis represents the node degrees (log10) and the y-axis the frequency of nodes having that degree (log10).

With regard to bicliques, 2,202 bicliques were detected in the E. coli network, with small bicliques being most prevalent and with c/p biclique size c=4 and p=2 being the most frequent biclique (106 times in the input network). The analyzed human network contained 22,879 bicliques. Here, larger bicliques were more frequent relative to *E. coli* and bicliques of size c=4 and p=2 being most frequent with 419 occurrences (Figure 4).

**Figure 4.**
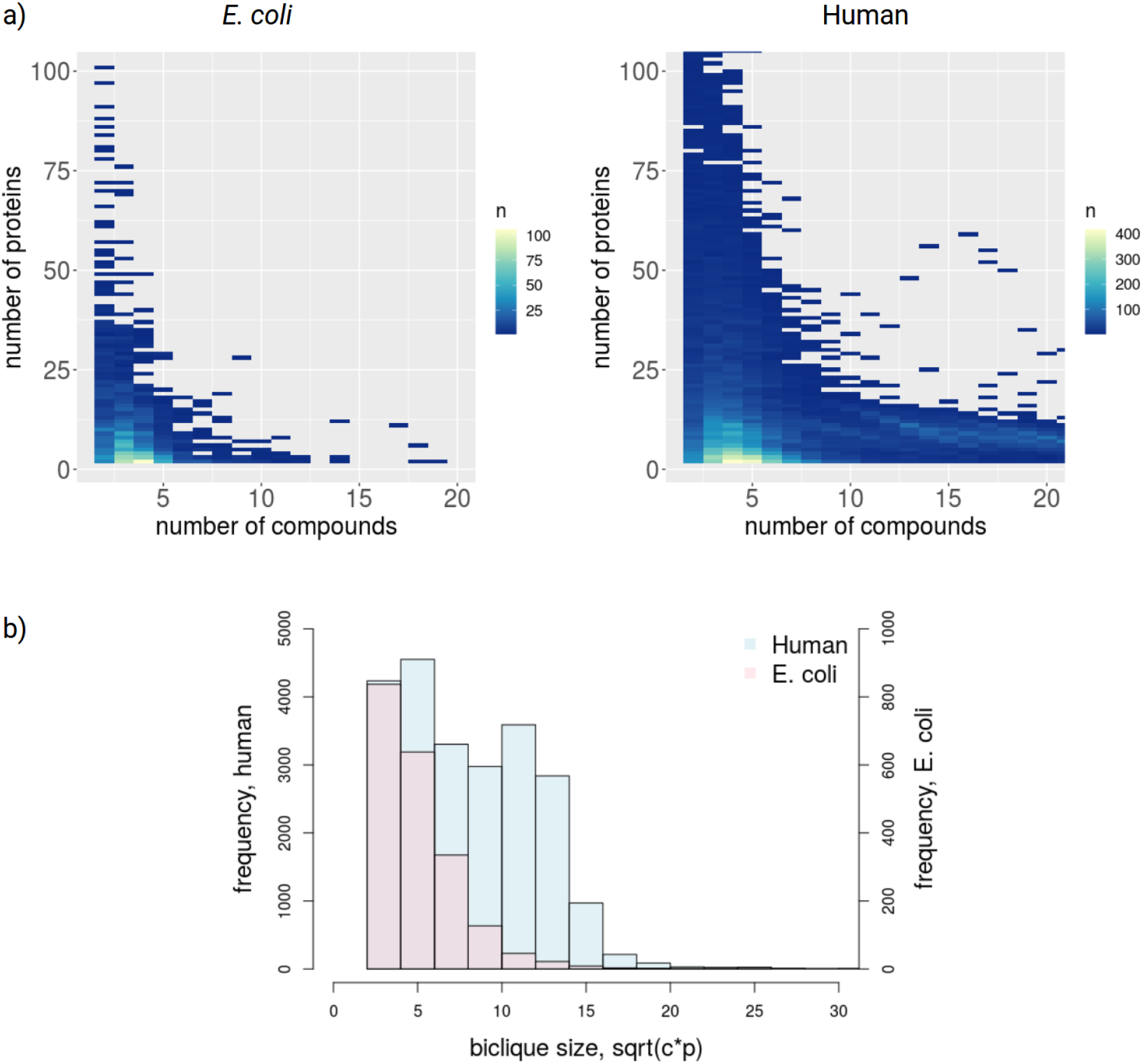
Size-dependent biclique frequency. a) Left: E. coli heatmap for all compound/ protein biclique sizes up to 100 proteins and 20 compounds. Bicliques including four compounds and two proteins are the most common with n = 106. (Total number of bicliques n = 2,022) a) Right: human heatmap for all c/p biclique sizes up to 100 proteins and 20 compounds. Bicliques including 4 compounds and 2 proteins are the most common with n = 419. (Total number of bicliques n = 22,879). b) Comparison of frequencies of bicliques of different sizes captured as a single number to allow for better comparison of E. coli vs. human and defined as sqrt(c*p), where c is the number of compounds and p the number of proteins (histogram clipped at size sqrt(c*p) = 30).

#### 3.1.1 Similarity of compounds and proteins

In bicliques, by definition, all member-compounds interact with the same set of proteins, and likewise, all member-poteins interact with the same set of compounds. As this agreement with regard to their respective molecular binding partners must have a molecular basis, it seems reasonable to hypothesize that proteins and compounds that are part of the same biclique are structurally similar. Indeed, proteins belonging to the same biclique show significantly higher sequence similarity (p<2.2E-16, with sequence similarity used as a proxy of structural similarity) than found between proteins that belong to different bicliques (Figure 5). Likewise, compounds that are members of the same biclique display greater chemical similarity than compounds that are not members of the same bicliques (p<2.2E-16, Figure 5). Interestingly, the difference of similarity within or across bicliques seems stronger for proteins (Cohen’s d effect size = 1.55) than for compounds (Cohen’s d=0.49). Assuming that the difference of similarity scale does not affect effect size, this may reflect that proteins may have several binding sites (e.g. for substrate and co-factors) such that with regard to compounds, diversity is greater than for proteins.

**Figure 5.**
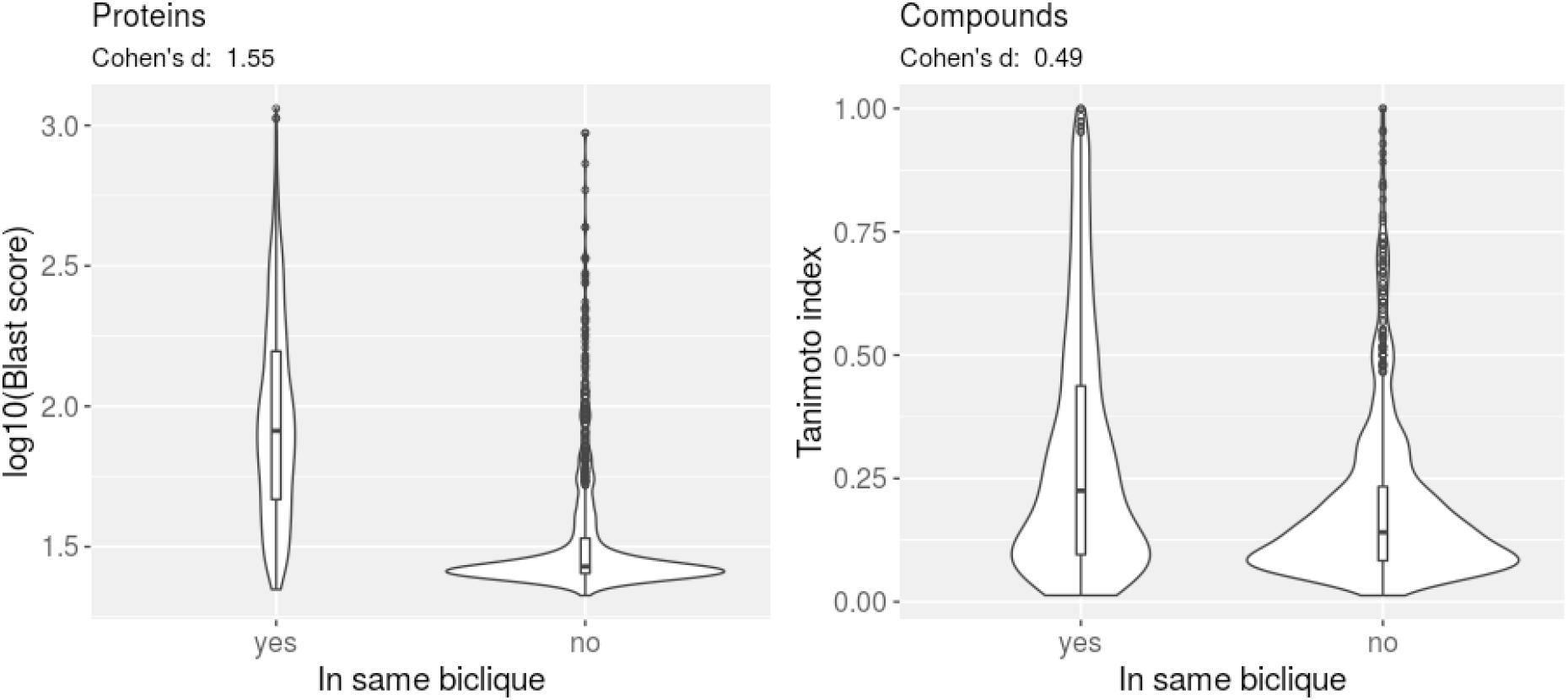
Molecular similarity of compounds and proteins within the same and between different bicliques. Violin plots of molecular similarity measures (blastp-score for proteins, Tanimoto index for compounds, see Materials and Methods) of compounds and proteins that are part of the same biclique or not with Cohen’s d effect sizes indicated, respectively. Distributions are based on 1000 randomly selected within and across-different molecule pairs. Corresponding Wilcoxon rank sum test p-values: compounds < 2.2e-16, proteins < 2.2e-16.

### 3.2 Biclique extension, Prediction results

Based on the obtained reference networks, we employed the logic of biclique extension as laid out in Figure 1 to predict novel interactions between compounds and proteins.

#### 3.2.1 E. coli, Compound-Protein-Interactions (CPI)

First, we tested the performance of the biclique extension method on the *E. coli* CPI reference network. Performed in a cross-validation-type test setting, we calculated true-positive rates (TPRs) and false-positive rates (FPRs) to evaluate our prediction results. In each of the 10 performed test runs, randomly chosen 10 percent of true interactions were considered unknown for the purpose of prediction, allowing to compute mean rates and associated standard deviations on the respective hold-out set.

The number of predicted interactions strongly depended on the sizes of bicliques with tested size-thresholds ranging from two to eight for the minimum number of nodes on either side of the biclique (nodes representing compounds (c) and proteins (p)), from 87 to approximately 171,154 interaction candidates (Figure 6). Note that biclique size refers to biclique size thresholds, which means that we defined the minimum size of bicliques, which were considered for calculation of performance measures. For example, by applying a threshold of c= 5 and p=2, all interactions were predicted using bicliques of this size and, in addition, all occurring bicliques of larger size, with higher or equal number of compounds (c) and proteins (p) such as c=6 and p=2 were also included.

**Figure 6.**
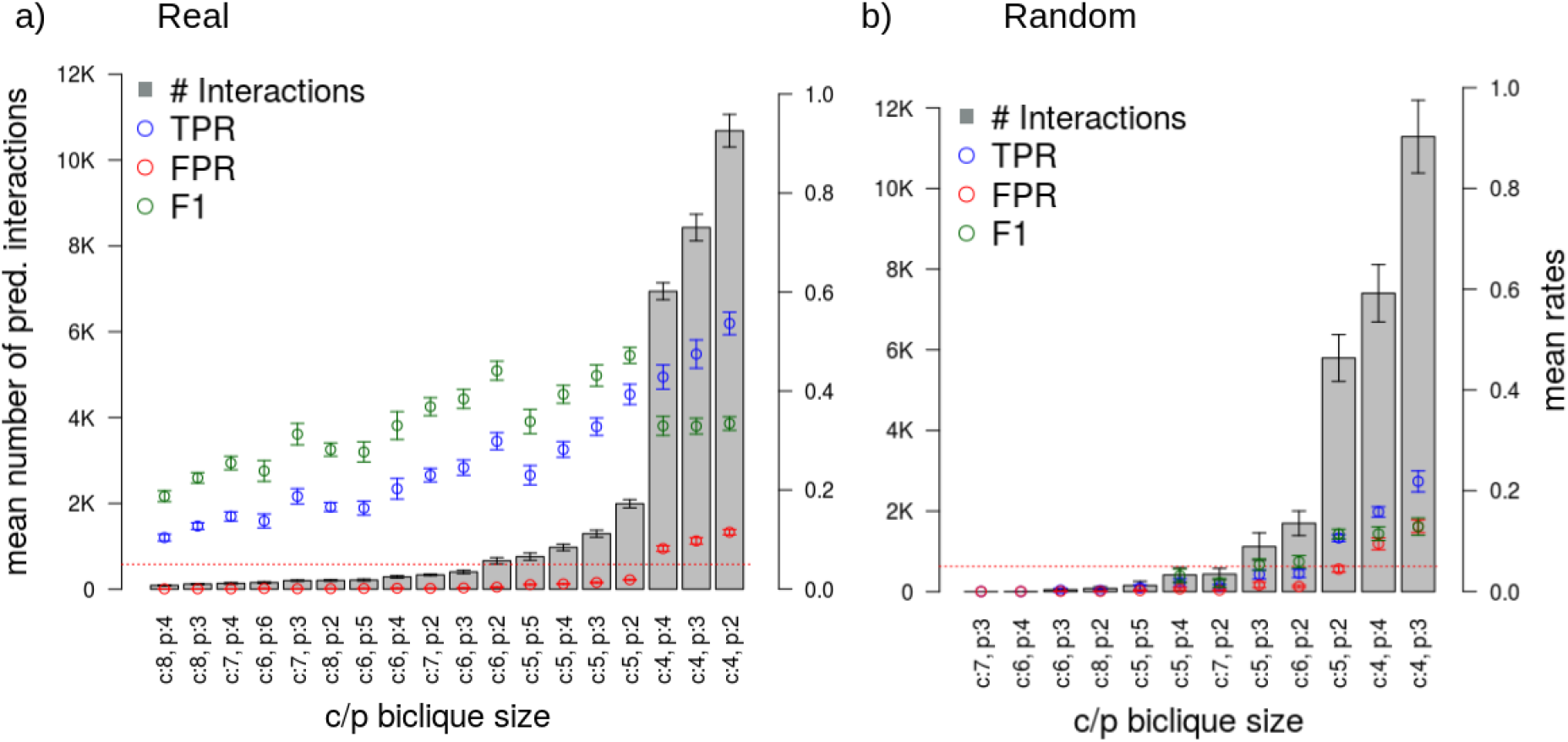
Results of the biclique-extension-based interaction predictions using a confidence threshold of 0.4 and with ten repetitions on different random samples (10% removed true interactions).On the left-hand side, the results for the real network are shown (a) and on the right-hand side the results for the randomized network (b). Sorting of data corresponding to different biclique sizes in ascending order of the number of predicted interactions (grey bars, average of 10 runs). The best obtained biclique size was c = 5 and p = 2, with maximal TPR with concurrent TPR<5% and highest F1-score. The red dotted line marks the 5%-line to allow for better visual clarity with regard to FPR. Note that the shown c/p biclique sizes represent the subset of all possible biclique sizes resulting in less than 12,000 predicted interactions, i.e. about twice the number of interactions in the input reference network. (Supplementary Tables 3, 4). Error bars correspond to standard deviations.

Generally, smaller maximal biclique sizes resulted in more predicted interactions, as the predicted interactions for larger bicliques were also included, and also, due to their increased occurrence in comparison to larger bicliques (Figure 4). The highest F1 measure associated also with the highest fraction of true-positive interactions with a FPR below 5% out of all predicted interactions was obtained by using c/p-biclique size-thresholds of five compounds (c=5) and two proteins (p=2) (Figure 6a). Applying these c/p-biclique size parameters resulted in an average TPR of 39.32% (Figure 6a, Supplementary Table 3). To test whether the biclique extension method proves both sensitive and specific and exploits actual biological information as present in the used reference network, we compared the obtained TPRs and FPRs with corresponding rates obtained after randomization of the network using the edge switch algorithm. As expected, for randomized data, we obtained significantly lower TPRs in comparison to real data, for example TPR of 10.62% for c=5, p=2 (Figure 6b, Supplementary Table 4). By contrast, the FPRs observed in real network data and random data were found at similar levels.

As some compounds, in particular those that act as cofactors such as ATP or NADPH etc. bind to many proteins - those compounds are often dubbed “currency metabolites”, we checked how removing them (for a complete list of compounds considered “currency metabolites’’, see Supplementary table 14), impacts the prediction performance. While the number of predicted interactions dropped significantly (from 1,994 to 457 for biclique size c:5, p:2), which is to be expected, the prediction performance was affected only slightly (F1 scores 47.2 vs. 40.2) (Figure 6a, Supplementary Figure 5 and Supplementary Table 15). Thus, the reported prediction performance does not only rely on high-degree interactors.

We also tested the performance when allowing the addition of two edges to declare a compound or protein to be a member of the corresponding biclique. Here, we considered larger bicliques only (minimum number of four compounds or proteins on the corresponding side), to better balance added vs. pre-existing interactions. As expected, this resulted in an increased number of predicted interactions for these larger bicliques (Supplementary Figure 5) as well as increased the FPRs. The prediction performance was comparable with a maximum F1 score of 42.14 and could not be increased compared to allowing only one edge addition.

We aimed to identify the prediction parameters that would be effective in reducing the number of experimental tests needed to obtain a set of validated novel interactions. Our prediction results with maximum F1-score, maximum sensitivity (TPR) and associated FPRs below 5% resulted in a confirmation rate (i.e. ratio of true-positive interactions to the number of predicted interactions) of at least 12% (Table 2). In comparison to the broad scale experimental studies with confirmation rates of about 5% (Piazza et al. 2018), this substantially increases the rate of validated interactions. Relying on larger bicliques would significantly increase the expected confirmation rate (71.43% for c:8, p:4), but at the expense of a much smaller number of predictions and true-positives (Table 2).

**Table 2.**
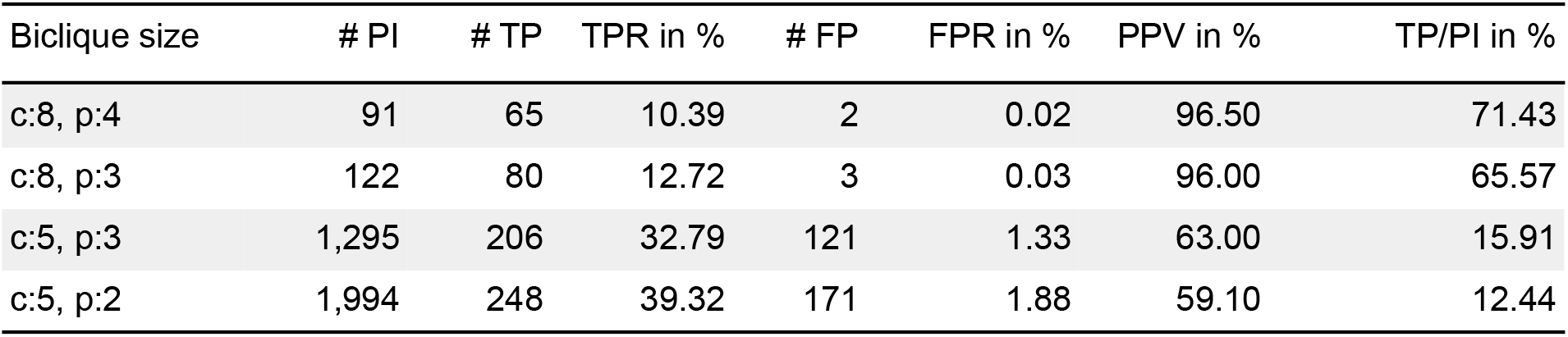
Biclique prediction performance with different minimum number of compounds (c) and proteins (p). The number of edges represents the number of predicted interactions. PI - predicted interactions, TP/PI provides an estimation of the expected validation success when tested experimentally. Chosen data correspond to bicliques with associated mean FPRs below 5%, n_repeats_=10. PPV - positive predictive value. Two large bicliques (c:8,p:4/3) and two middle sized bicliques (c:5, p:3/2) are listed, including the biclique size which was identified as the optimum (c:5, p2). For a complete data table, see Supplementary table 3.

#### 3.2.2 Human CPI

We applied the biclique extension method to the larger human compound protein interaction network, which included 44,322 interactions (confidence threshold of 0.4 and after filtering).

In ten performed validation runs, we used a set of five percent randomly drawn true interactions of the input network as positive controls, which were removed from the network prior to prediction, and a downloaded set of negative interactions as true negatives (see Methods). In dependence of the sizes of thresholds ranging from two to nine for the minimum number of compounds and proteins on either side of the biclique (maximum c/p-biclique size), 1,648 to approximately 1.6 million candidate interactions were predicted (Figure 7a, Supplementary Table 7). The TPRs showed an increase with decreasing biclique size, while the FPRs were below 5% for all bicliques with more than four compounds and two proteins (Figure 7a, Supplementary Table 7). The highest F1 and, thus, the best biclique size threshold was obtained for c=4, p=2 (F1 = 76.72%), followed by c=5, p=2 (F1 = 75.99%), which were also found to perform best in *E. coli* (Figure 6a). We also tested the insertion of two edges for larger bicliques. This showed no major effect on the mean FPRs and the mean TPRs (Supplementary Figure 4). For a randomized input network, the number of predicted interactions increased significantly compared to the real network (Figure 7b and d). The c/p-biclique size c=4, p=2 yielded a mean number of 201,212 predicted interactions in the real network and 983,616 interactions in the randomized network (Supplementary table 7 and 8). As expected, TPRs decreased, FPRs increased, and F1s decreased accordingly in the randomized network and consistently across all biclique threshold sizes (Figure 7b vs 7a).

**Figure 7.**
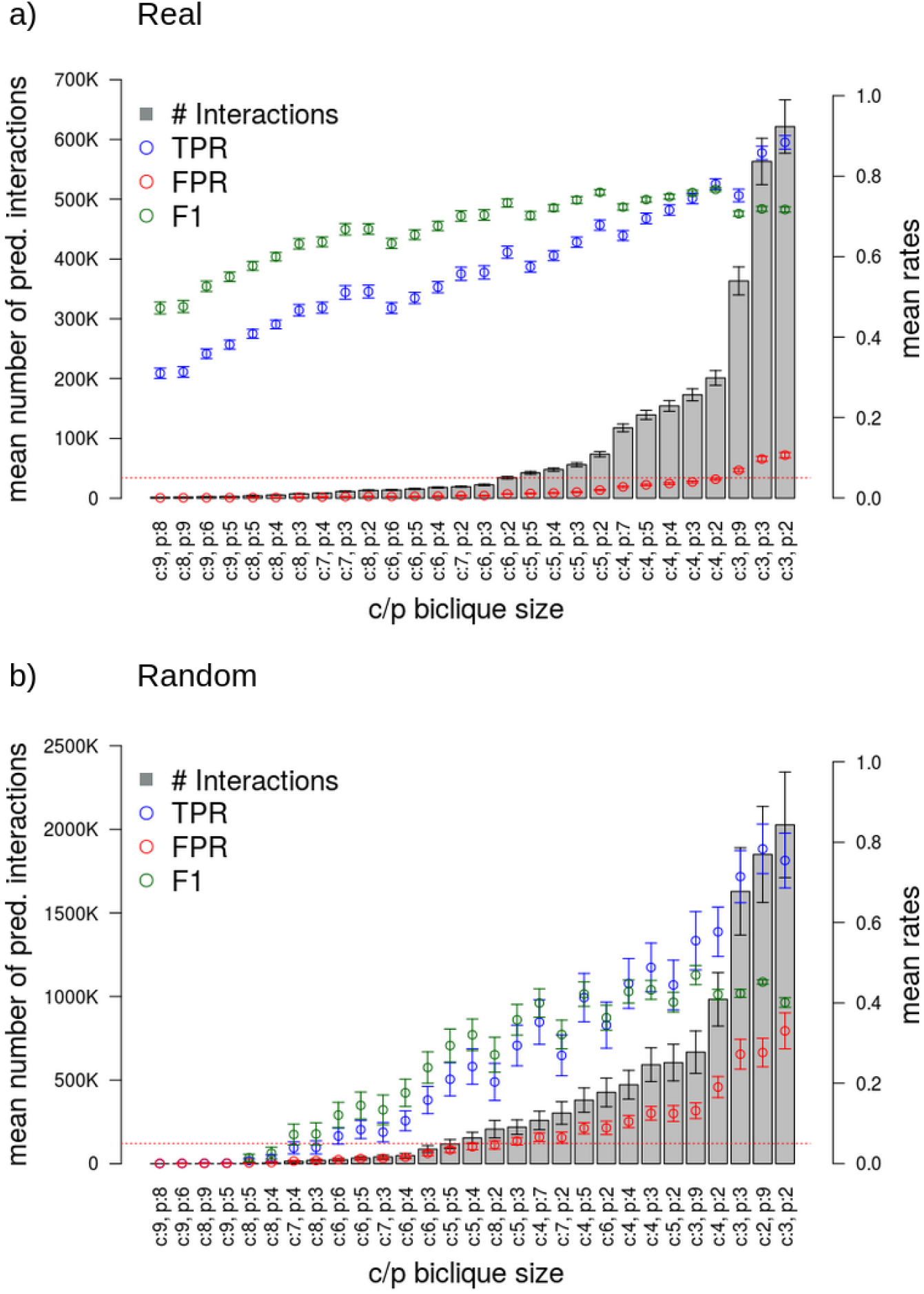
Results of the biclique-extension-based interaction predictions using a confidence threshold of 0.4 and with ten repetitions on different random samples (5% removed true interactions). Sorting of data corresponding to different biclique sizes in ascending order of the number of predicted interactions (grey bars, average of 10 runs). The best obtained biclique size was c = 4 and p = 2, with maximal TPR with concurrent TPR<5% and highest F1-score. The red dotted line marks the 5%-line to allow for better visual clarity with regard to FPR. Note that the shown c/p biclique sizes represent the subset of all possible biclique sizes resulting in less than 700,000 predicted interactions for real data and comparable biclique size ranges for random data. (Supplementary Tables 7, 8). Error bars correspond to standard deviations.

### 3.3 Dependency on the choice of interaction confidence threshold

Following our investigations of identifying the optimal c/p-biclique size and generating comparable random data, we also tested the dependence of our results on the confidence threshold used to create the reference network based on the STITCH *E.coli* interaction data. We compared the true-positive rates (TPR) and false-positive rates (FPR) in relation to the number of predicted interactions as a representation of c/p-biclique size (each point corresponds to a specific biclique size) using a range of confidence thresholds from 0.4 to 0.6.

In general, we obtained similar TPRs and FPRs for each tested confidence threshold (Figure 8). The only small differences indicate that interactions predicted by our biclique method are not strongly dependent on the underlying levels of confidence in the analyzed interaction network. Among the three tested thresholds, confidence scores 0.5 and 0.4 (green and black line) yielded the highest TPRs and the lowest FPRs, while the network with highest confidence score 0.6, surprisingly, performed worse, possibly explained by the smaller networks, and thus reduced information in a biclique sense, associated with more stringent thresholds.

**Figure 8.**
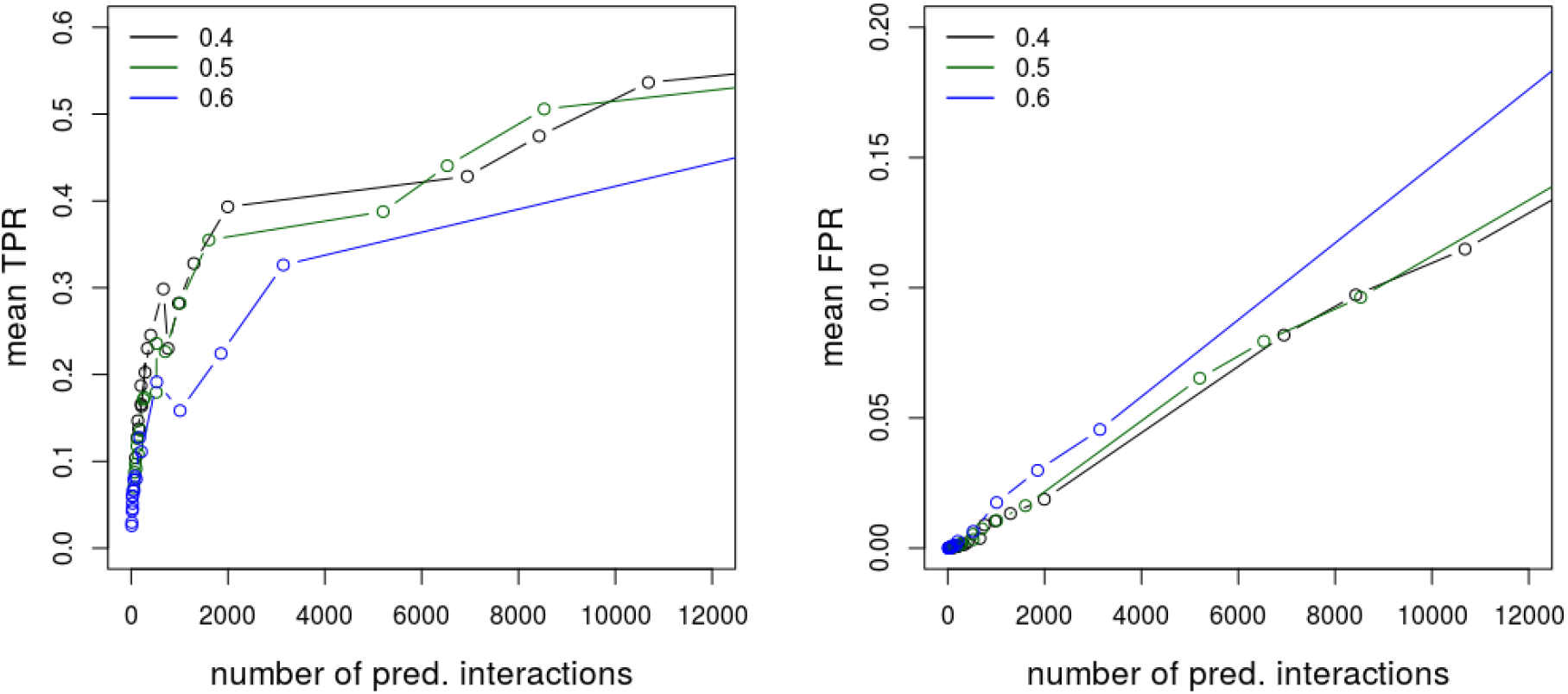
Mean TPRs (left) and mean FPRs (right) in dependence on the number of predicted interactions in the E. coli interaction network (data from 10 repeat runs of different random sample sets). Colors represent the three different confidence thresholds applied to the STITCH network to create the initial basis network. Every circle corresponds to a particular biclique size (c/p) as listed in Figure 6. Note the different scales of the y-axes.

To determine optimal parameter settings, we also inspected the precision-recall statistic for the *E. coli* networks based on different STITCH confidence scores and maximum biclique sizes. Consistent with our findings reported in Figure 8, we obtained the best results applying confidence thresholds of 0.4 and 0.5 (Figure 9). As expected, larger maximal biclique sizes resulted in higher precision. Here, the biclique-based prediction rests on more support, because a larger number of interactions is known in these bicliques already rendering the logic of biclique extension more applicable. And that logic states that if a compound binds to n-1 other proteins that another set of m compounds bind to, then the one missing interaction for that compound likely occurs as well. And with larger n and m, this logic is more compelling and for structural reasons (Figure 5). The same holds for proteins and their interactions to compounds.

**Figure 9.**
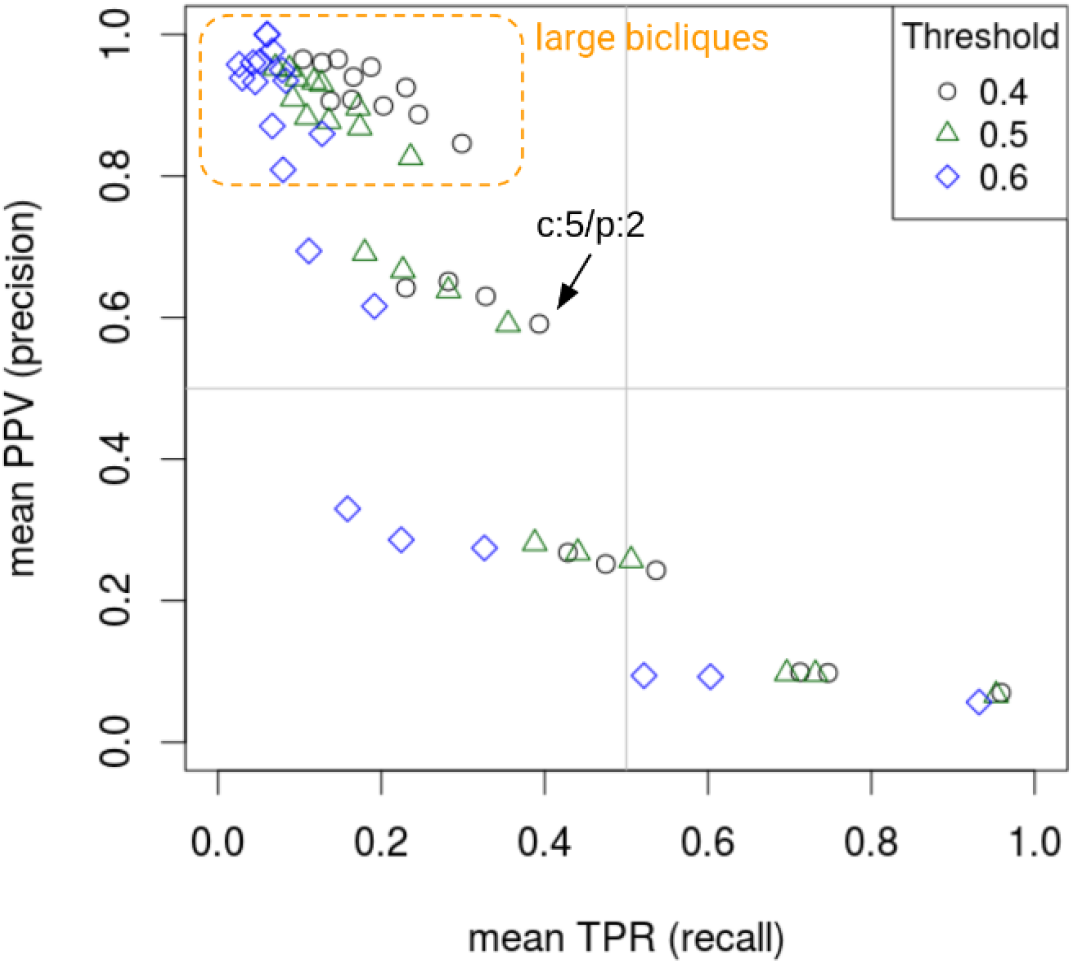
The mean true-positive rate (TPR or “recall”) in relation to the mean positive predictive value (PPV or “precision”) calculated for our prediction results using three different applied confidence thresholds (data from 10 repeat runs of different random sample sets). In the best case, the points are in the upper right corner, maximizing both precision and recall. Here, the effect of many more negatives (~ 9,000) than positives (~ 600), combined with imperfect predictions prevents reaching higher precision and recall. Large bicliques with c=6-8 yielded high precision but low recall. The found optimal (highest F1 score) biclique size (c:5, p:2) is indicated in the graph. There are only slight differences between the PPV and TPR of networks based on different STITCH confidence thresholds, especially for the 0.4 and 0.5 thresholds.

### 3.4 KEGG Enrichment Analysis on the *E. coli* network

Following performance evaluation, suggesting high prediction power, we applied the biclique extension method to the whole *E. coli* STITCH network as input. Applying a confidence input score of 0.4 and setting the biclique-size threshold to the one with detected highest F1 score (c:5,p:2), our method predicts 2,666 novel interactions between 127 compounds and 444 proteins. Of note, 681 (25.5%) of those interactions were in fact already contained in the STITCH network, but with an experimental confidence level below our threshold. (The complete list of predictions is available as a Supplementary File).

To inspect our prediction results with regard to biological function (proteins) and their biochemical processes (compounds), we performed a KEGG annotation enrichment analysis on the *E. coli* network to discern metabolic pathways that are enriched in the input STITCH network, in bicliques, and in the predicted interactions. The input list of proteins was compared to the KEGG database as a reference to determine pathway enrichment.

In the complete filtered STITCH network of known physical interactions, amino acid metabolism pathways showed the highest fold-enrichment (’Phenylalanine, tyrosine and tryptophan biosynthesis’ and ‘Alanine, aspartate and glutamate metabolism’, Figure 10a) relative to the KEGG annotation. Furthermore, other central metabolic processes (e.g. ‘Glycolysis’), but also the ‘biosynthesis of secondary metabolites’ were found overrepresented, with the latter being associated with the highest significance (smallest adjusted p-value. We obtained almost identical results for the set of proteins that were members of at least one biclique (not shown), as almost all proteins were detected to be part of a biclique (all but 10).

**Figure 10.**
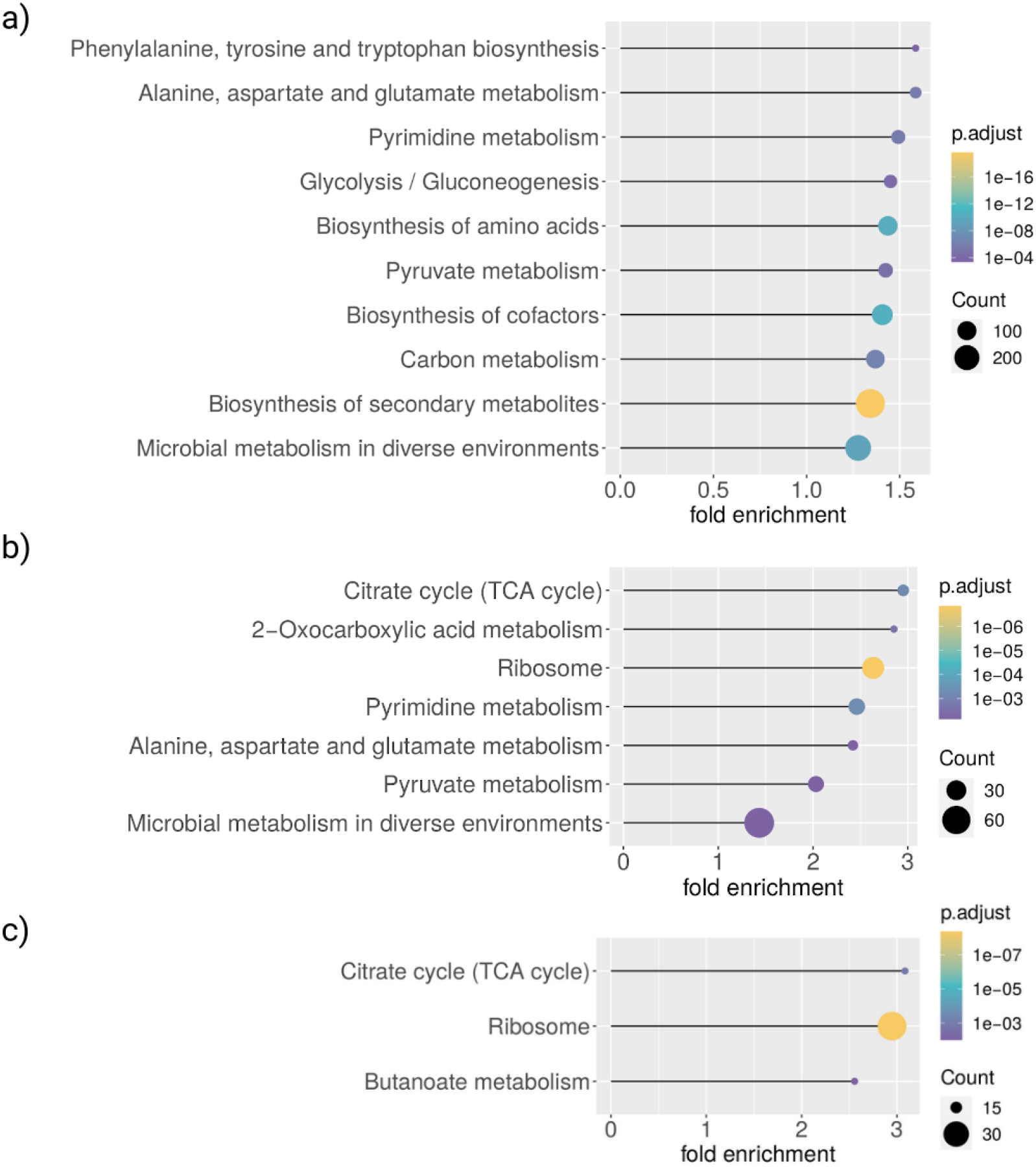
KEGG annotation enrichment analysis of proteins in the E. coli network using the R package clusterProfiler. Shown are up to 10 KEGG-categories with p.adjust<0.01, ordered by fold enrichment. Control for multiple testing via the Benjamini-Hochberg method (default in clusterProfiler). a) Enriched KEGG pathways for all proteins of the input network, b) for proteins which are in bicliques up to c/p-biclique size of c=5 and p=2 (size with highest F1 score) and c) for proteins of predicted interactions applying the same maximum c/p-biclique size. For a complete table see Supplementary Tables 11-13.

The evaluation of our predictions had revealed an optimal maximum c/p-biclique size of c=5 and p=2. Thus, we additionally analyzed the enrichment in this subset of proteins that belong to bicliques of this optimal maximal size. Here, ‘TCA Cycle’ showed the largest fold-enrichment (Figure 10b), followed by ‘2-Oxocarboxylic acid metabolism’ and ‘ribosome’. All three were not reported for the full interaction set (Figure 10a vs 10b). The next four categories reported enriched in bicliques (‘Pyrimidine-’, ‘Alanine, aspirated and glutamate-’, ‘Pyruvate-’ metabolism, and ‘Microbial metabolism in diverse environments’) were already reported overrepresented in the whole input network. Thus, with regard to biochemical processes, bicliques show both characteristic as well as common process association.

Next, we analyzed the enrichment for the predicted interactions. These predictions were also calculated based on the bicliques with the maximum c/p-biqlique size c=5 and p=2. As we predicted new interactions which are connected to the existing bicliques of the input network, we obtained a similar enrichment as for the input bicliques (overlap of two out of three), with one more process ‘Butanoate metabolism’ reported significant. The largest enriched categories were ‘TCA Cycle’ and ‘Ribosome’ (Figure 10c). 75 of 420 proteins were only included in the predicted interactions but not part of existing bicliques of the applied c/p-biclique size.

## 4. DISCUSSION

Aiming to contribute to a deeper understanding of the function of compounds, proteins, and their interactions in metabolic and cellular networks, we searched for the missing links in compound-protein (CPI) networks with an explicit focus on metabolite-protein interactions. We applied the method of biclique extension, which works by discovering incomplete bicliques in a given network and postulating all edges missing for completion of these bicliques as potential novel connections. Based only on network topology, we predicted novel interactions in an *E. coli* and in a human compound-protein interaction network, assuming that predicted edges represent candidates of novel interactions. Bipartite extension makes use of known interactions between compounds and proteins to complete the interaction network. Each compound or protein that gets connected to an existing biclique by insertion of only one or two edges was considered to represent a novel interaction partner. As such, bipartite extension can be viewed to rely on little input information: no detailed knowledge and prediction of binding modes, energetics, and biochemical process involvement is necessary. At the same time, it only works with sufficient knowledge of existing interactions, and thus, requires rather extensive prior information. Nonetheless, we believe the biclique extension method to represent an alternative approach to molecular docking approaches that require molecular structural information and a precise description in interaction potentials and other machine learning methods (Chen et al. 2016; Tsubaki, Tomii, and Sese 2019;Tsubaki, Tomii, and Sese 2010). As we showed, despite bipartite extension not imposing any molecular information, it does implicitly capture molecular similarity as a determining factor for the validity of inference of interaction (Figure 5). We believe the primary application scenario of this method lies in the completion of networks. By this study, we have demonstrated that bipartite-based methods hold great potential when focusing on metabolite-protein interactions in addition to the demonstrations of their successful applications to drug-target interaction prediction and drug-repositioning (Lotfi Shahreza et al. 2018).

We achieved a sensitivity of 39.32% for our predictions of compound-protein interactions on *E. coli*, while keeping the number of false positives below 5%, which resulted in an associated precision of 59.1% (Figure 6a, Supplementary Table 3). We demonstrated an even better prediction performance for the human network, in which we obtained a maximum sensitivity of nearly 78.05%, increasing with the total number of predicted interactions (Figure 7, Supplementary Table 7). In both species, prediction performance levels were significantly and substantially above random predictions. We found that our performance rates depended on the chosen c/p-biclique sizes. The best sensitivity for both networks was obtained when including smaller c/p-biclique sizes, even though larger bicliques occur at higher frequency in the human network than in the *E. coli* network. The larger we chose the c/p-biclique sizes, the larger was the set of compounds and proteins with known interactions captured by these bicliques. As is to be expected, this resulted in an increase of sensitivity and precision as well as a decrease of FPRs, as the logic of the biclique extension rests on more support (Figure 6, 7). However, large c/p-biclique sizes also lead to a lower number of bicliques in the query network, and, thus, result in a lower total number of predictable interactions (Supplementary tables 3,4,7,8). For the *E. coli* and the human network, we found the optimal minimum c/p-biclique sizes were of four or five compounds and two proteins, with regard to maximum TPRs and minimum FPRs. Consequently, this seems to be an optimal minimum c/p-biclique size. Nevertheless, different c/p-biclique sizes should be tested on the input network. Another effective filter criterion would be to limit the total number of predicted interactions, as this also keeps the FPRs low. Consistently, we found best performance for bicliques with more compounds than proteins in them (Figures 6, 7). As this holds also for the frequencies of biclique sizes for the input network (Figure 4), we assume this to be purely a statistical effect.

Evidently, bipartite extension depends on the input network to be correct. While this can be assumed to be true for the used STITCH network - within the limits of the employed confidence assessment - we also found that the biclique extension proved robust with regard to the chosen STITCH confidence scores (Figures 8, 9). This may reveal a strength of the biclique extension method. It integrates over many interactions to make predictions and may thus prove error tolerant.

We compared our obtained performance rates for the *E. coli* network to rates from a recent experimental study (*E. coli* CPIs, Piazza et al. 2018) to evaluate whether our approach would be able to improve the prediction of novel compound-protein interactions in general. Indeed, focusing tests on predicted interactions obtained by the biclique extension method would increase the fraction of validated interactions in relation to the number of tests. In detail, Piazza et al. tested 34,186 interactions experimentally and reported 1,719 validated interactions, out of which 1,487 were novel targets. This reflects a validation rate of 4.35% for novel interactions and 5.03% relative to all validated interactions (including also previously known interactions). Our predictions resulted in mean validation rates of 12.44% with TPRs of 39.32% up to validation rates of 71.43% with TPRs of 10.39%, relative to the number of predicted interactions and with FPRs below 5% (n=10). Noticeably, we can even assume these rates to actually be higher, as several of the predicted interactions might contain additional true positive interactions that are not yet part of the set of real positive cases found by experimental testing. Clearly, Piazza et al. went for systematic testing and did not aim to optimize for the highest rate of validated interactions. However, achieving an increase of validated interactions by simultaneously reducing the number of required tests would save time and resources. Consequently, novel interaction partners predicted by biclique extension can be highly supportive when considered for experimental testing.

With regard to biochemical process involvement of compounds and proteins represented in bicliques and biclique-based interaction predictions, we found TCA-cycle, ribosome overrepresented in both (Figure 10 b,c). As the TCA-cycle is a central biochemical integration hub with relatively tight metabolites-enzyme interactions described before (albeit in yeast, Durek and Walther 2008) the appearance of this process in this statistic seems very plausible. Furthermore, amongst TCA-enzymes/proteins themselves, many interactions have been reported (in *Arabidopsis thaliana*: Youjun Zhang et al. 2018) rendering an associated compound-protein biclique interactions more likely, with the notion of support of substrate channeling referred to as metabolons observed for TCA-cycle. Of particular interest here are the reported new interactions within the TCA-cycle (see Supplementary File) that await experimental verification. By contrast, as the ribosome is not immediately considered associated with metabolism, but with translation, its appearance seems surprising. However, as the ribosome is a multiprotein complex with many proteins binding to co-factors such as ATP and ADP, a densely knit interaction network with associated predictions can be explained as well. Of note also, even when probing the whole input STITCH interaction network relative to the KEGG annotation, pronounced overrepresentation of a number of metabolic functions have been observed (Figure 10a). Thus, physical interactions form a much denser network than that of biochemical reaction based substrate/product - enzyme interactions captured by KEGG.

In summary, we demonstrated that biclique extension is indeed helpful for the prediction of CPIs in naturally occurring metabolic and cellular networks. Bipartite extension works under minimal assumptions that are solely inferred from observed interaction networks and their topology and is not limited by the currently known biochemical and physico-chemical determinants of compound-protein interactions, which offers a high potential to find novel interaction candidates and to support efficient experimental testing. The biclique extension methodology can be readily applied to all species with available CPI interactions (STITCH). With this study, we provide a basis that allows choosing the parameters of biclique-extension-based predictions and provide expected performance levels of their applications in the context of metabolite-protein interaction networks.

## Acknowledgements

We wish to thank Zoran Nikoloski, Joachim Kopka, and Detlef Groth for helpful discussions, suggestions, and insightful comments.

## Notes

### Competing Interest Statement

The authors have declared no competing interest.

