## Supplementary File for "Biclique extension as an effective approach to predict novel interaction partners in metabolic compound-protein interaction networks"

Supplementary information

### Supplementary Figures

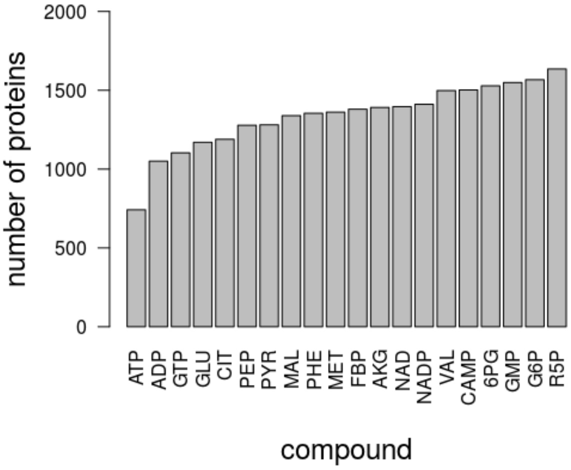


***Supplementary Figure 1***. *Reported number of negative interactions (no interaction observed) for the 20 compounds contained in the Piazza.negatives (Supplementary table 1).*

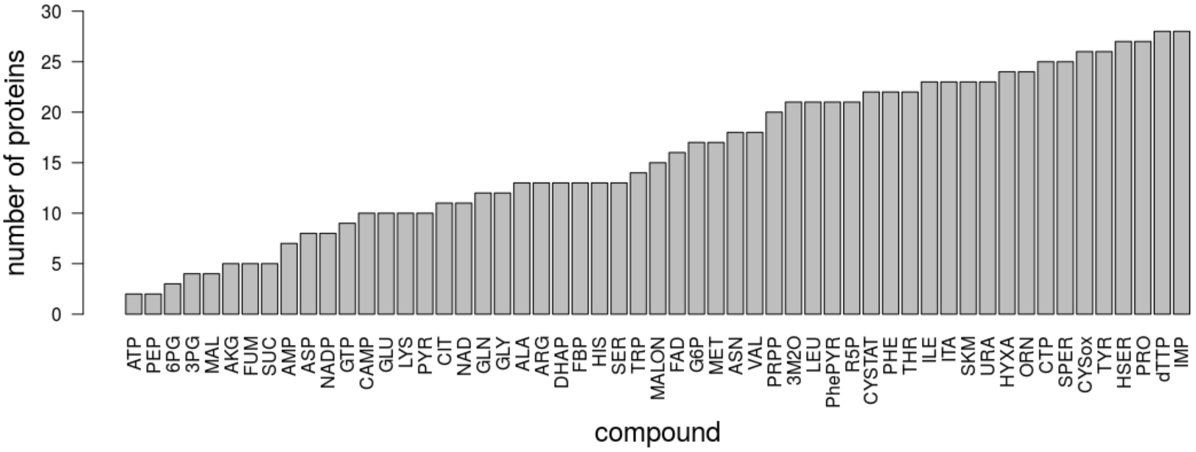


***Supplementary Figure 2***. *Reported number of negative interactions (no interaction observed) for the 55 compounds contained in the Diether.negatives (Supplementary table 2).*

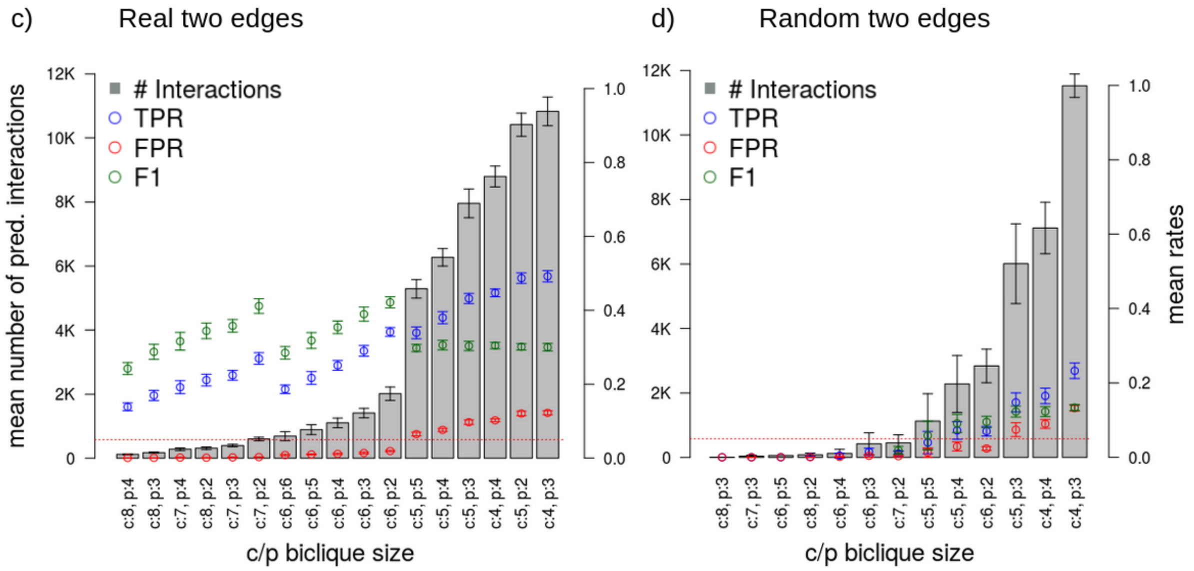


***Supplementary Figure 3.*** *Results of the biclique-extension-based interaction predictions using a confidence threshold of 0.4 and with ten repetitions on different random samples (10% removed true interactions). With insertion of one edge (a,b) and insertion of two edes (c,d) for larger bicliques (minimum of four nodes on the side to which interactions are added). On the left-hand side, the results for the real network are shown (a,c) and on the right-hand side the results for the randomized network (b,d). Sorting of data corresponding to different biclique sizes in ascending order of the number of predicted interactions (grey bars, average of 10 runs). The best obtained biclique size under the one-edge addition scenaria was c = 5 and p = 2, with maximal TPR with concurrent TPR<5% and highest F1-score. For the two-edge addition mode, c=6, p=2 proved optimal. The red dotted line marks the 5%-line to allow for better visual clarity with regard to FPR. Note that the shown c/p biclique sizes represent the subset of all possible biclique sizes resulting in less than 12,000 predicted interactions, i.e. about twice the number of interactions in the input reference network. (Supplementary Tables 5, 6). Error bars correspond to standard deviations.*

c) Real two edges

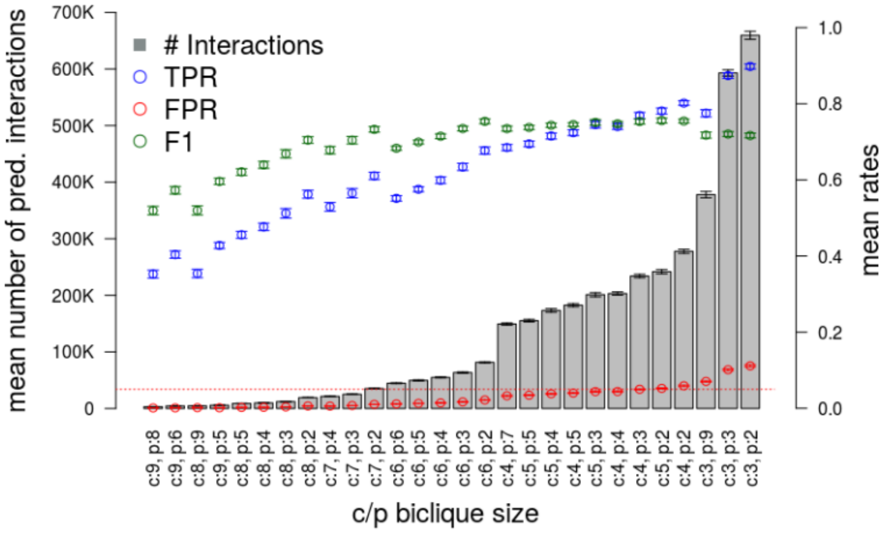


d) Random two edges

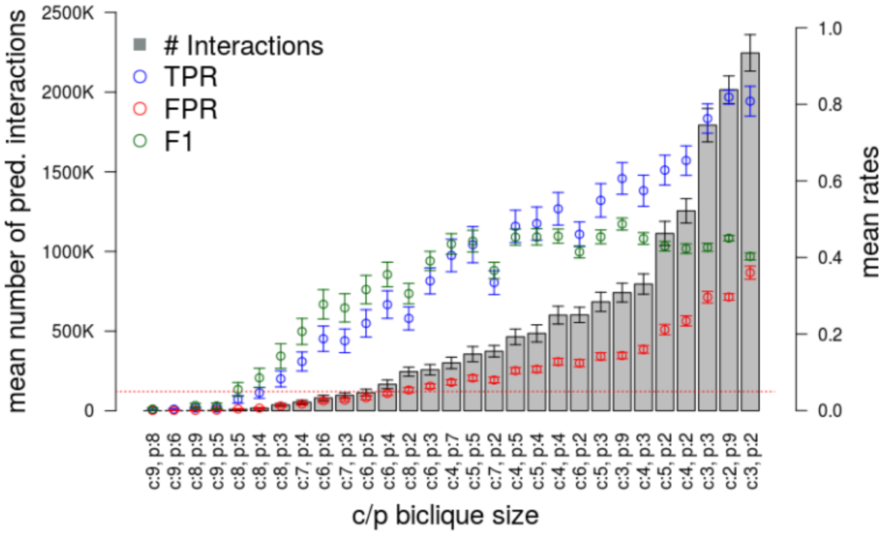


***Supplementary Figure 4.*** *Results of the biclique-extension-based interaction predictions using a confidence threshold of 0.4 and with ten repetitions on different random samples (5% removed true interactions). With insertion of two edes for larger bicliques (minimum of four nodes on the side to which interactions are added). Sorting of data corresponding to different biclique sizes in ascending order of the number of predicted interactions (grey bars, average of 10 runs). The best obtained biclique size under the two-edge addition mode was c=6, p=2, with highest F1. The red dotted line marks the 5%-line to allow for better visual clarity with regard to FPR. Note that the shown c/p biclique sizes represent the subset of all possible biclique sizes resulting in less than 700,000 predicted interactions for real data and comparable biclique size ranges for random data. (Supplementary Table 9-10). Error bars correspond to standard deviations*

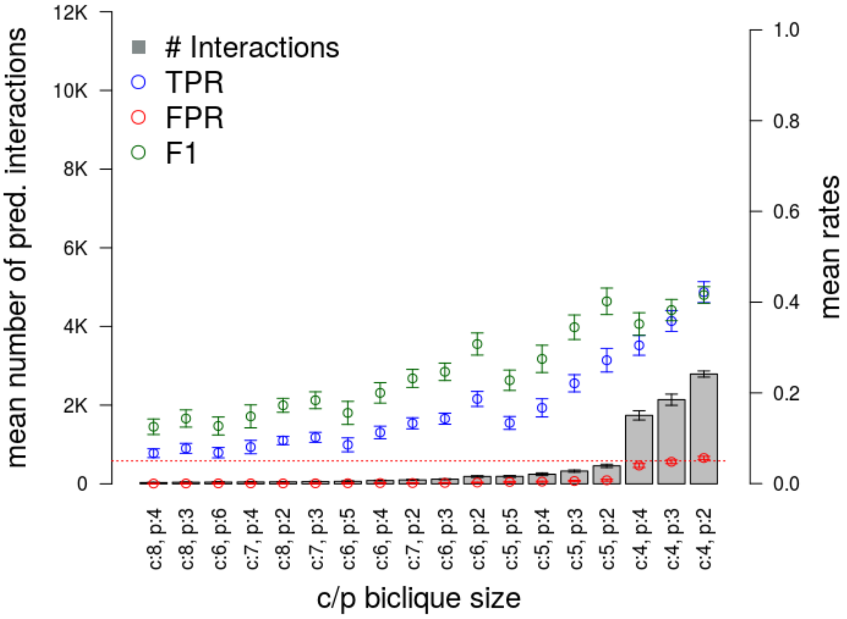


***Supplementary Figure 5.*** *Results of biclique-extension-based interaction prediction for E. coli after removing currency metabolites (see Supplementary table 9) from the network and using a confidence threshold of 0.4 and with ten repetitions on different random samples. Insertion of one edge was allowed for predictions. Sorting of data corresponding to different biclique sizes in ascending order of the number of predicted interactions (grey bars, average of 10 runs). The best obtained biclique size under the one-edge addition scenaria was c = 4/5 and p = 2, with maximal TPR with concurrent TPR<5%. Note that the shown c/p biclique sizes represent the subset of all possible biclique sizes resulting in less than 12,000 predicted interactions, i.e. about twice the number of interactions in the input reference network.*

### Supplementary Tables

***Supplementary Table 1****: List of all 20 compounds included in the validation dataset of Piazza et al (Piazza et al. 2018).*

| Abbreviation | Name | KEGG ID |
|---|---|---|
| ATP | Adenosine triphosphate | C00002 |
| NAD | Nicotinamide adenine dinucleotide | C00003 |
| NADP | Nicotinamide adenine dinucleotide phosphate | C00006 |
| ADP | Adenosine diphosphate | C00008 |
| Pyr | Pyruvate | C00022 |
| L-Gln | L-Glutamate | C00025 |
| AKG | 2-Oxoglutarate | C00026 |
| GTP | Guanosine triphosphate | C00044 |
| L-Met | L-Methionine | C00073 |
| PEP | Phosphoenolpyruvate | C00074 |
| L-Phe | L-Phenylalanine | C00079 |
| G6P | D-Glucose 6-phosphate | C00092 |
| R5P | alpha-D-Ribose 5-phosphate | C00117 |
| GMP | Guanosine monophosphate | C00144 |
| Mal | L-Malate | C00149 |
| Cit | Citrate | C00158 |
| L-Val | L-Valine | C00183 |
| PGP | 6-Phospho-D-gluconate | C00345 |
| FBP | D-Fructose 1,6-bisphosphate | C00354 |
| cAMP | 3',5'-cyclic adenosine monophosphate | C00575 |

***Supplementary Table 2**: List of all  compounds included in the validation dataset of Diether et al (Diether et al. 2019).*

| Abbreviation | Name | KEGG ID |
|---|---|---|
| 3M2O | 2-Ketoisovalerate, 2-Oxoisovalerate | C00141 |
| AKG | Ketoglutarate | C00026 |
| THR | Threonine | C00188 |
| G6P | alpha-D-glucose 6-phosphate | C00092 |
| AMP | Adenosine monophosphate | C00020 |
| ARG | Arginine | C00062 |
| ASN | Asparagine | C00152 |
| ATP | MgATP | C00002 |
| CAMP | cAMP | C00575 |
| CIT | Citrate | C00158 |
| ORN | L-Ornithine | C00077 |
| 6PG | 6-Phosphogluconic acid | C00345 |
| ASP | Aspartate | C00049 |
| MAL | L-Malate | C00149 |
| CTP | ara-CTP | C00063 |
| CYSox | Cysteine | C00097 |
| ALA | Alanine | C00041 |
| GLU | Glutamate | C00025 |
| PRO | Proline | C00148 |
| SER | D-Serine | C00065 |
| DHAP | Dihydroxyacetone phosphate | C00111 |
| FAD | Flavin adenine dinucleotide | C00016 |
| FBP | Fructose-1,6-diphosphate | C00354 |
| FUM | Fumarate | C00122 |
| 3PG | 3-Phosphoglycerate | C00197 |
| GLN | Glutamine | C00064 |
| GLY | Glycine | C00037 |
| GTP | Guanosine triphosphate | C00044 |
| HIS | Histidine | C00135 |

| | | |
|---|---|---|
| HSER | L-Homoserine | C00263 |
| HYXA | Hypoxanthine | C00262 |
| ILE | Isoleucine | C00407 |
| IMP | Inosine monophosphate | C00130 |
| ITA | Itaconate | C00490 |
| CYSTAT | L-Cystathionine | C02291 |
| LEU | Leucine | C00123 |
| LYS | Lysine | C00047 |
| MALON | Malonate | C00383 |
| MET | Methionine | C00073 |
| NAD | Diphosphopyridine nucleotide | C00003 |
| NADP | Nicotinamide adenine dinucleotide phosphate | C00006 |
| PHE | Phenylalanine | C00079 |
| PhePYR | Phenylpyruvate | C00166 |
| PEP | Phosphoenolpyruvate | C00074 |
| PRPP | 5-Phosphoribosyl-1-pyrophosphate | C00119 |
| PYR | Pyruvate | C00022 |
| SKM | Shikimic acid | C00493 |
| SPER | Spermidine | C00315 |
| SUC | Succinate | C00042 |
| TRP | Tryptophan | C00078 |
| dTTP | Thymidine 5'- | C00459 |
| TYR | Tyrosine | C00082 |
| URA | Uracil | C00106 |
| VAL | L-Valine | C00183 |
| R5P | Riboside 5-monophosphate | C00117 |

***Supplementary Table 3.*** *In relation to Figure 6a. Statistical results of biclique calculations for E. coli with different minimum number of compounds (c) and proteins (p) on each side of the biclique using a confidence threshold of 0.4 and allowing insertion of one edge to complete a biclique. The number of edges represents the number of predicted interactions. Datasets are sorted according to the number of edges. All values are the mean values of n=10 repetitions. Note that the underlying reference data was unbalanced with a mean number of positives n= 630 and a mean number of negatives n = 9,116. Dataset with maximum F1 score referred to in the text is highlighted.*

| Dataset | # Edges | # TP | TPR in % | # FP | FPR in % | PPV in % | F1 in % |
|---|---|---|---|---|---|---|---|
| Biclique c:8, p:4 | 91 | 65 | 10.39 | 2 | 0.02 | 96.50 | 18.75 |
| Biclique c:8, p:3 | 122 | 80 | 12.72 | 3 | 0.03 | 96.00 | 22.46 |
| Biclique c:7, p:4 | 137 | 92 | 14.68 | 4 | 0.04 | 96.50 | 25.46 |
| Biclique c:6, p:6 | 157 | 87 | 13.75 | 9 | 0.10 | 90.60 | 23.86 |
| Biclique c:7, p:3 | 202 | 118 | 18.73 | 6 | 0.06 | 95.40 | 31.28 |
| Biclique c:8, p:2 | 205 | 104 | 16.58 | 7 | 0.07 | 94.00 | 28.18 |
| Biclique c:6, p:5 | 214 | 103 | 16.36 | 10 | 0.12 | 90.80 | 27.70 |
| Biclique c:6, p:4 | 288 | 128 | 20.25 | 14 | 0.16 | 89.90 | 33.02 |
| Biclique c:7, p:2 | 335 | 145 | 23.01 | 12 | 0.13 | 92.50 | 36.84 |
| Biclique c:6, p:3 | 404 | 154 | 24.54 | 20 | 0.22 | 88.70 | 38.42 |
| Biclique c:6, p:2 | 664 | 188 | 29.86 | 34 | 0.38 | 84.60 | 44.11 |
| Biclique c:5, p:5 | 761 | 145 | 23.00 | 81 | 0.89 | 64.20 | 33.83 |
| Biclique c:5, p:4 | 977 | 178 | 28.21 | 95 | 1.04 | 65.10 | 39.34 |
| Biclique c:5, p:3 | 1,295 | 206 | 32.79 | 121 | 1.33 | 63.00 | 43.12 |
| **Biclique c:5, p:2** | **1,994** | **248** | **39.32** | **171** | **1.88** | **59.10** | **47.20** |
| Biclique c:4, p:4 | 6,943 | 270 | 42.83 | 746 | 8.18 | 26.80 | 32.95 |
| Biclique c:4, p:3 | 8,428 | 299 | 47.47 | 887 | 9.73 | 25.20 | 32.90 |
| Biclique c:4, p:2 | 10,683 | 338 | 53.65 | 1,047 | 11.48 | 24.30 | 33.44 |
| Biclique c:3, p:3 | 42,962 | 449 | 71.27 | 4,142 | 45.44 | 9.90 | 17.38 |
| Biclique c:3, p:2 | 48,039 | 470 | 74.69 | 4,342 | 47.62 | 9.80 | 17.32 |
| Biclique c:2, p:2 | 136,643 | 604 | 95.89 | 8,119 | 89.06 | 7.00 | 13.05 |

***Supplementary Table 4.*** *In relation to Figure 6b. Statistical results of biclique calculations for E. coli randomized network with different minimum number of compounds (c) and proteins (p) on each side of the biclique using a confidence threshold of 0.4 and allowing insertion of one edge. The number of edges represents the number of predicted interactions. Datasets are sorted according to the number of edges. All values are the mean values of n=10 repetitions. Note that the underlying reference data was unbalanced with a mean number of positives n= 503 and a mean number of negatives n = 8,586.*

| Dataset | # Edges | # TP | TPR in % | # FP | FPR in % | PPV in % | F1 in % |
|---|---|---|---|---|---|---|---|
| Biclique c:7, p:3 | 2 | 0 | 0.00 | 0 | 0.00 | NaN | NaN |
| Biclique c:6, p:4 | 5 | 0 | 0.00 | 1 | 0.01 | NaN | NaN |
| Biclique c:6, p:3 | 50 | 1 | 0.26 | 5 | 0.06 | NaN | NaN |
| Biclique c:8, p:2 | 83 | 1 | 0.22 | 4 | 0.05 | 12.97 | NaN |
| Biclique c:5, p:5 | 156 | 4 | 0.74 | 18 | 0.21 | NaN | NaN |
| Biclique c:5, p:4 | 417 | 9 | 1.75 | 44 | 0.52 | 17.78 | 3.11 |
| Biclique c:7, p:2 | 438 | 5 | 0.91 | 23 | 0.26 | 16.02 | 1.72 |
| Biclique c:5, p:3 | 1,120 | 17 | 3.38 | 111 | 1.29 | 14.10 | 5.33 |
| Biclique c:6, p:2 | 1,698 | 18 | 3.58 | 85 | 1.00 | 17.84 | 5.92 |
| Biclique c:5, p:2 | 5,796 | 53 | 10.62 | 382 | 4.44 | 12.44 | 11.42 |
| Biclique c:4, p:4 | 7,402 | 79 | 15.80 | 813 | 9.47 | 9.03 | 11.46 |
| Biclique c:4, p:3 | 11,875 | 108 | 21.54 | 1,174 | 13.67 | 8.55 | 12.21 |
| Biclique c:4, p:2 | 21,210 | 153 | 30.46 | 1,657 | 19.30 | 8.47 | 13.25 |
| Biclique c:3, p:3 | 61,568 | 303 | 60.26 | 4,953 | 57.69 | 5.76 | 10.52 |
| Biclique c:3, p:2 | 74,844 | 321 | 63.93 | 5,278 | 61.47 | 5.74 | 10.53 |
| Biclique c:2, p:2 | 158,768 | 466 | 92.74 | 8,035 | 93.58 | 5.48 | 10.35 |

***Supplementary Table 5.*** *In relation to Supplementary Figure 3c. Statistical results of biclique calculations for E. coli with different minimum number of compounds (c) and proteins (p) on each side of the biclique using a confidence threshold of 0.4 and allowing insertion of two edges. The number of edges represents the number of predicted interactions. Datasets are sorted according to the number of edges. All values are the mean values of n=10 repetitions. Note that the underlying reference data was unbalanced with a mean number of positives n= 624 and a mean number of negatives n = 9,089. Dataset with maximum F1 score referred to in the text is highlighted.*

| Dataset | # Edges | # TP | TPR in % | # FP | FPR in % | PPV in % | F1 in % |
|---|---|---|---|---|---|---|---|
| Biclique c:8, p:4 | 122 | 86 | 13.86 | 3 | 0.03 | 96.40 | 24.22 |
| Biclique c:8, p:3 | 175 | 106 | 16.95 | 6 | 0.06 | 95.00 | 28.75 |
| Biclique c:7, p:4 | 283 | 120 | 19.19 | 13 | 0.14 | 90.10 | 31.61 |
| Biclique c:8, p:2 | 315 | 132 | 21.11 | 9 | 0.10 | 93.60 | 34.43 |
| Biclique c:7, p:3 | 398 | 140 | 22.41 | 17 | 0.19 | 89.10 | 35.79 |
| Biclique c:7, p:2 | 602 | 168 | 26.94 | 25 | 0.27 | 87.30 | 41.16 |
| Biclique c:6, p:6 | 690 | 116 | 18.62 | 76 | 0.83 | 60.90 | 28.49 |
| Biclique c:6, p:5 | 894 | 136 | 21.70 | 91 | 1.00 | 59.80 | 31.82 |
| Biclique c:6, p:4 | 1,104 | 157 | 25.08 | 103 | 1.14 | 60.30 | 35.40 |
| Biclique c:6, p:3 | 1,412 | 181 | 29.00 | 126 | 1.38 | 59.40 | 38.95 |
| **Biclique c:6, p:2** | **2,015** | **213** | **34.13** | **174** | **1.91** | **55.10** | **42.14** |
| Biclique c:5, p:5 | 5,290 | 212 | 33.92 | 590 | 6.50 | 26.60 | 29.79 |
| Biclique c:5, p:4 | 6,272 | 237 | 38.02 | 694 | 7.63 | 25.60 | 30.58 |
| Biclique c:5, p:3 | 7,956 | 270 | 43.20 | 883 | 9.71 | 23.40 | 30.34 |
| Biclique c:4, p:4 | 8,796 | 279 | 44.72 | 930 | 10.23 | 23.10 | 30.45 |
| Biclique c:5, p:2 | 10,414 | 304 | 48.71 | 1,097 | 12.06 | 21.80 | 30.11 |
| Biclique c:4, p:3 | 10,828 | 307 | 49.20 | 1,112 | 12.24 | 21.60 | 30.01 |
| Biclique c:4, p:2 | 13,877 | 347 | 55.53 | 1,326 | 14.59 | 20.80 | 30.25 |
| Biclique c:3, p:3 | 43,529 | 447 | 71.55 | 4,096 | 45.07 | 9.90 | 17.39 |
| Biclique c:3, p:2 | 48,783 | 472 | 75.54 | 4,369 | 48.07 | 9.90 | 17.50 |
| Biclique c:2, p:2 | 137,270 | 600 | 96.09 | 8,099 | 89.11 | 7.00 | 13.05 |

***Supplementary Table 6.*** *In relation to Supplementary Figure 3d. Statistical results of biclique calculations for E. coli randomized network with different minimum number of compounds (c) and proteins (p) on each side of the biclique using a confidence threshold of 0.4 and allowing insertion of one edge. The number of edges represents the number of predicted interactions. Datasets are sorted according to the number of edges. All values are the mean values of n=10 repetitions. Note that the underlying reference data was unbalanced with mean number of positives n= 508 and the mean number of negatives n = 8,585.*

| Dataset | # Edges | # TP | TPR in % | # FP | FPR in % | PPV in % | F1 in % |
|---|---|---|---|---|---|---|---|
| Biclique c:8, p:3 | 10 | 0 | 0.00 | 0 | 0.00 | NaN | NaN |
| Biclique c:7, p:3 | 40 | 0 | 0.00 | 2 | 0.02 | NaN | NaN |
| Biclique c:6, p:5 | 60 | 0 | 0.00 | 3 | 0.03 | 0.00 | NaN |
| Biclique c:8, p:2 | 82 | 1 | 0.12 | 3 | 0.04 | NaN | NaN |
| Biclique c:6, p:4 | 125 | 2 | 0.43 | 8 | 0.10 | 27.90 | NaN |
| Biclique c:6, p:3 | 424 | 7 | 1.45 | 32 | 0.37 | 20.14 | NaN |
| Biclique c:7, p:2 | 453 | 5 | 1.00 | 26 | 0.30 | 16.84 | 1.87 |
| Biclique c:5, p:5 | 1,127 | 20 | 3.96 | 108 | 1.26 | 17.53 | 5.84 |
| Biclique c:5, p:4 | 2,280 | 37 | 7.21 | 251 | 2.92 | 13.40 | 9.10 |
| Biclique c:6, p:2 | 2,839 | 36 | 7.00 | 203 | 2.36 | 15.12 | 9.50 |
| Biclique c:5, p:3 | 6,009 | 75 | 14.76 | 638 | 7.43 | 10.75 | 12.29 |
| Biclique c:4, p:4 | 7,116 | 84 | 16.49 | 772 | 9.00 | 9.84 | 12.29 |
| Biclique c:4, p:3 | 12,264 | 121 | 23.76 | 1,225 | 14.27 | 9.02 | 13.06 |
| Biclique c:5, p:2 | 17,234 | 141 | 27.69 | 1,471 | 17.13 | 8.75 | 13.29 |
| Biclique c:4, p:2 | 24,173 | 173 | 33.93 | 1,910 | 22.25 | 8.29 | 13.33 |
| Biclique c:3, p:3 | 61,511 | 310 | 60.97 | 5,040 | 58.70 | 5.79 | 10.58 |
| Biclique c:3, p:2 | 76,023 | 330 | 64.96 | 5,377 | 62.63 | 5.79 | 10.63 |
| Biclique c:2, p:2 | 159,408 | 471 | 92.64 | 8,061 | 93.89 | 5.52 | 10.42 |

***Supplementary Table 7.*** *In relation to Figure 7a. Statistical results of biclique calculations for human with different minimum number of compounds (c) and proteins (p) on each side of the biclique using a confidence threshold of 0.4 and allowing insertion of one edge. The number of edges represents the number of predicted interactions. Datasets are sorted according to the number of edges. Dataset with maximum F1 score referred to in the text is highlighted. Note that the underlying reference data was unbalanced with mean number of positives n= 2,099 and the mean number of negatives n = 11,458.*

| Dataset | # Edges | # TP | TPR in % | # FP | FPR in % | PPV in % | F1 in % |
|---|---|---|---|---|---|---|---|
| Biclique c:9, p:8 | 1,648 | 651 | 31.01 | 6 | 0.05 | 99.13 | 47.23 |
| Biclique c:8, p:9 | 1,928 | 658 | 31.34 | 7 | 0.06 | 98.99 | 47.59 |
| Biclique c:9, p:6 | 2,447 | 752 | 35.84 | 8 | 0.07 | 99.00 | 52.62 |
| Biclique c:9, p:5 | 2,990 | 800 | 38.12 | 10 | 0.09 | 98.77 | 55.00 |
| Biclique c:8, p:5 | 3,904 | 857 | 40.82 | 13 | 0.11 | 98.51 | 57.71 |
| Biclique c:8, p:4 | 5,225 | 906 | 43.15 | 15 | 0.13 | 98.42 | 59.99 |
| Biclique c:8, p:3 | 7,257 | 980 | 46.67 | 21 | 0.19 | 97.86 | 63.18 |
| Biclique c:7, p:4 | 8,442 | 993 | 47.33 | 28 | 0.25 | 97.24 | 63.66 |
| Biclique c:7, p:3 | 11,234 | 1,073 | 51.12 | 39 | 0.34 | 96.53 | 66.82 |
| Biclique c:8, p:2 | 13,115 | 1,077 | 51.33 | 44 | 0.38 | 96.09 | 66.90 |
| Biclique c:6, p:6 | 13,667 | 991 | 47.23 | 42 | 0.37 | 95.94 | 63.29 |
| Biclique c:6, p:5 | 15,348 | 1,044 | 49.73 | 47 | 0.41 | 95.73 | 65.44 |
| Biclique c:6, p:4 | 17,936 | 1,100 | 52.41 | 52 | 0.46 | 95.46 | 67.66 |
| Biclique c:7, p:2 | 19,160 | 1,170 | 55.74 | 68 | 0.59 | 94.51 | 70.11 |
| Biclique c:6, p:3 | 22,291 | 1,177 | 56.09 | 70 | 0.61 | 94.41 | 70.36 |
| Biclique c:6, p:2 | 34,180 | 1,282 | 61.09 | 114 | 1.00 | 91.83 | 73.36 |
| Biclique c:5, p:5 | 42,367 | 1,207 | 57.48 | 130 | 1.14 | 90.26 | 70.23 |
| Biclique c:5, p:4 | 47,868 | 1,265 | 60.26 | 144 | 1.25 | 89.81 | 72.12 |
| Biclique c:5, p:3 | 55,987 | 1,335 | 63.58 | 169 | 1.48 | 88.77 | 74.09 |
| Biclique c:5, p:2 | 73,593 | 1,424 | 67.86 | 225 | 1.97 | 86.37 | 75.99 |
| Biclique c:4, p:7 | 117,610 | 1,369 | 65.24 | 318 | 2.78 | 81.16 | 72.33 |
| Biclique c:4, p:5 | 139,361 | 1,459 | 69.50 | 375 | 3.27 | 79.57 | 74.18 |
| Biclique c:4, p:4 | 154,342 | 1,503 | 71.61 | 412 | 3.60 | 78.50 | 74.88 |
| Biclique c:4, p:3 | 172,985 | 1,564 | 74.49 | 456 | 3.98 | 77.44 | 75.93 |
| **Biclique c:4, p:2** | **201,212** | **1,638** | **78.05** | **534** | **4.66** | **75.45** | **76.72** |

| | | | | | | | |
|---|---|---|---|---|---|---|---|
| Biclique c:3, p:9 | 363,364 | 1,579 | 75.21 | 790 | 6.89 | 66.69 | 70.68 |
| Biclique c:3, p:3 | 563,164 | 1,801 | 85.80 | 1,112 | 9.71 | 61.86 | 71.87 |
| Biclique c:3, p:2 | 621,522 | 1,856 | 88.41 | 1,221 | 10.65 | 60.36 | 71.72 |
| Biclique c:2, p:9 | 982,566 | 1,811 | 86.26 | 1,717 | 14.99 | 51.36 | 64.37 |
| Biclique c:2, p:2 | 1,592,215 | 2,028 | 96.63 | 2,557 | 22.32 | 44.28 | 60.71 |

***Supplementary Table 8.*** *In relation to Figure 7b. Statistical results of biclique calculations for human randomized networks with different minimum number of compounds (c) and proteins (p) on each side of the biclique using a confidence threshold of 0.4 and allowing insertion of one edge. The number of edges represents the number of predicted interactions. Datasets are sorted according to the number of edges. Dataset with maximum F1 score referred to in the text is highlighted. Note that the underlying reference data was unbalanced with mean number of positives n= 1,888 and the mean number of negatives n = 11,501.*

| Dataset | # Edges | # TP | TPR in % | # FP | FPR in % | PPV in % | F1 in % |
|---|---|---|---|---|---|---|---|
| Biclique c:9, p:8 | 31 | 0 | 0.00 | 1 | 0.01 | NaN | NaN |
| Biclique c:9, p:6 | 95 | 1 | 0.03 | 1 | 0.01 | NaN | NaN |
| Biclique c:8, p:9 | 153 | 1 | 0.08 | 2 | 0.02 | NaN | NaN |
| Biclique c:9, p:5 | 274 | 2 | 0.10 | 3 | 0.03 | NaN | NaN |
| Biclique c:8, p:5 | 1,628 | 13 | 0.71 | 14 | 0.13 | 45.58 | 1.39 |
| Biclique c:8, p:4 | 4,061 | 25 | 1.34 | 27 | 0.24 | 46.42 | 2.58 |
| Biclique c:7, p:4 | 14,262 | 74 | 3.91 | 69 | 0.60 | 51.08 | 7.22 |
| Biclique c:8, p:3 | 19,309 | 76 | 4.02 | 83 | 0.72 | 46.87 | 7.34 |
| Biclique c:6, p:6 | 23,600 | 129 | 6.84 | 108 | 0.94 | 54.42 | 12.06 |
| Biclique c:6, p:5 | 33,118 | 159 | 8.45 | 140 | 1.22 | 53.10 | 14.47 |
| Biclique c:7, p:3 | 39,876 | 147 | 7.80 | 148 | 1.29 | 49.47 | 13.36 |
| Biclique c:6, p:4 | 47,983 | 201 | 10.66 | 182 | 1.58 | 52.55 | 17.60 |
| Biclique c:6, p:3 | 87,188 | 298 | 15.79 | 291 | 2.53 | 50.73 | 23.89 |
| Biclique c:5, p:5 | 117,156 | 395 | 20.98 | 393 | 3.42 | 50.21 | 29.37 |
| Biclique c:5, p:4 | 154,501 | 455 | 24.15 | 482 | 4.19 | 48.62 | 32.04 |
| Biclique c:8, p:2 | 206,770 | 383 | 20.31 | 533 | 4.63 | 41.91 | 27.08 |
| Biclique c:5, p:3 | 218,890 | 554 | 29.38 | 639 | 5.56 | 46.49 | 35.77 |
| Biclique c:4, p:7 | 258,861 | 664 | 35.20 | 754 | 6.56 | 46.89 | 39.96 |
| Biclique c:7, p:2 | 303,101 | 507 | 26.91 | 742 | 6.45 | 40.84 | 32.12 |

| | | | | | | | |
|---|---|---|---|---|---|---|---|
| Biclique c:4, p:5 | 380,442 | 779 | 41.31 | 1,010 | 8.78 | 43.62 | 42.20 |
| Biclique c:6, p:2 | 426,662 | 649 | 34.43 | 1,021 | 8.88 | 39.05 | 36.29 |
| Biclique c:4, p:4 | 472,403 | 845 | 44.83 | 1,201 | 10.44 | 41.37 | 42.81 |
| Biclique c:4, p:3 | 592,246 | 921 | 48.82 | 1,438 | 12.50 | 39.11 | 43.24 |
| Biclique c:5, p:2 | 604,874 | 838 | 44.45 | 1,433 | 12.46 | 37.04 | 40.17 |
| Biclique c:3, p:9 | 667,629 | 1,046 | 55.48 | 1,512 | 13.15 | 40.99 | 46.92 |
| Biclique c:4, p:2 | 983,616 | 1,088 | 57.69 | 2,190 | 19.04 | 33.32 | 42.07 |
| Biclique c:3, p:3 | 1,628,943 | 1,348 | 71.43 | 3,130 | 27.21 | 30.25 | 42.36 |
| Biclique c:2, p:9 | 1,849,606 | 1,478 | 78.33 | 3,178 | 27.63 | 31.89 | 45.19 |
| Biclique c:3, p:2 | 2,027,395 | 1,423 | 75.44 | 3,797 | 33.01 | 27.41 | 40.08 |
| Biclique c:2, p:2 | 4,392,612 | 1,725 | 91.39 | 6,587 | 57.27 | 20.86 | 33.91 |

***Supplementary Table 9.*** *In relation to Supplementary Figure 4c. Statistical results of biclique calculations for human with different minimum number of compounds (c) and proteins (p) on each side of the biclique using a confidence threshold of 0.4 and allowing insertion of two edges. The number of edges represents the number of predicted interactions. Datasets are sorted according to the number of edges. Dataset with maximum F1 score referred to in the text is highlighted. Note that the underlying reference data was unbalanced with mean number of positives n= 2,099 and the mean number of negatives n = 11,458.*

| Dataset | # Edges | # TP | TPR in % | # FP | FPR in % | PPV in % | F1 in % |
|---|---|---|---|---|---|---|---|
| Biclique c:9, p:8 | 3,204 | 769 | 36.65 | 15 | 0.13 | 98.09 | 53.36 |
| Biclique c:9, p:6 | 4,442 | 873 | 41.61 | 16 | 0.14 | 98.20 | 58.45 |
| Biclique c:8, p:9 | 4,739 | 766 | 36.51 | 23 | 0.20 | 97.08 | 53.06 |
| Biclique c:9, p:5 | 5,710 | 913 | 43.52 | 21 | 0.18 | 97.75 | 60.23 |
| Biclique c:8, p:5 | 8,631 | 966 | 46.04 | 31 | 0.27 | 96.89 | 62.42 |
| Biclique c:8, p:4 | 9,600 | 1,016 | 48.43 | 34 | 0.30 | 96.76 | 64.55 |
| Biclique c:8, p:3 | 11,897 | 1,096 | 52.24 | 46 | 0.40 | 95.97 | 67.65 |
| Biclique c:8, p:2 | 18,979 | 1,187 | 56.58 | 76 | 0.66 | 93.98 | 70.64 |
| Biclique c:7, p:4 | 20,732 | 1,140 | 54.34 | 63 | 0.55 | 94.76 | 69.07 |
| Biclique c:7, p:3 | 24,739 | 1,214 | 57.86 | 81 | 0.71 | 93.75 | 71.56 |
| Biclique c:7, p:2 | 34,242 | 1,304 | 62.15 | 114 | 0.99 | 91.96 | 74.17 |
| Biclique c:6, p:6 | 44,483 | 1,177 | 56.10 | 136 | 1.19 | 89.64 | 69.01 |
| Biclique c:6, p:5 | 48,985 | 1,221 | 58.20 | 151 | 1.32 | 88.99 | 70.38 |

| | | | | | | | |
|---|---|---|---|---|---|---|---|
| Biclique c:6, p:4 | 54,294 | 1,284 | 61.20 | 172 | 1.50 | 88.19 | 72.26 |
| Biclique c:6, p:3 | 62,067 | 1,359 | 64.78 | 200 | 1.74 | 87.17 | 74.33 |
| **Biclique c:6, p:2** | **80,384** | **1,443** | **68.78** | **249** | **2.17** | **85.28** | **76.15** |
| Biclique c:4, p:7 | 151,568 | 1,454 | 69.30 | 385 | 3.36 | 79.06 | 73.86 |
| Biclique c:5, p:5 | 158,559 | 1,478 | 70.45 | 403 | 3.51 | 78.58 | 74.29 |
| Biclique c:5, p:4 | 172,838 | 1,514 | 72.16 | 447 | 3.90 | 77.21 | 74.60 |
| Biclique c:4, p:5 | 184,450 | 1,536 | 73.21 | 470 | 4.10 | 76.57 | 74.85 |
| Biclique c:5, p:3 | 202,275 | 1,581 | 75.36 | 498 | 4.34 | 76.05 | 75.70 |
| Biclique c:4, p:4 | 204,738 | 1,570 | 74.83 | 505 | 4.40 | 75.66 | 75.24 |
| Biclique c:4, p:3 | 233,901 | 1,628 | 77.60 | 567 | 4.94 | 74.17 | 75.85 |
| Biclique c:5, p:2 | 242,027 | 1,646 | 78.46 | 601 | 5.24 | 73.25 | 75.77 |
| Biclique c:4, p:2 | 278,556 | 1,687 | 80.41 | 686 | 5.98 | 71.09 | 75.46 |
| Biclique c:3, p:9 | 379,548 | 1,631 | 77.74 | 831 | 7.24 | 66.25 | 71.53 |
| Biclique c:3, p:3 | 596,552 | 1,833 | 87.37 | 1,175 | 10.24 | 60.94 | 71.80 |
| Biclique c:3, p:2 | 668,174 | 1,885 | 89.85 | 1,299 | 11.32 | 59.20 | 71.38 |
| Biclique c:2, p:9 | 1,009,623 | 1,854 | 88.37 | 1,733 | 15.11 | 51.69 | 65.22 |
| Biclique c:2, p:2 | 1,631,265 | 2,046 | 97.52 | 2,613 | 22.78 | 43.92 | 60.56 |

***Supplementary Table 10.*** *In relation to Supplementary Figure 4d. Statistical results of biclique calculations for human randomized networks with different minimum number of compounds (c) and proteins (p) on each side of the biclique using a confidence threshold of 0.4 and allowing insertion of one edge. The number of edges represents the number of predicted interactions. Datasets are sorted according to the number of edges. Dataset with maximum F1 score referred to in the text is highlighted. Note that the underlying reference data was unbalanced with mean number of positives n= 1,880 and the mean number of negatives n = 11,506.*

| Dataset | # Edges | # TP | TPR in % | # FP | FPR in % | PPV in % | F1 in % |
|---|---|---|---|---|---|---|---|
| Biclique c:9, p:8 | 284 | 4 | 0.20 | 4 | 0.03 | 42.80 | 0.39 |
| Biclique c:9, p:6 | 749 | 7 | 0.36 | 8 | 0.07 | NaN | NaN |
| Biclique c:8, p:9 | 1,038 | 13 | 0.68 | 9 | 0.08 | 62.80 | 1.33 |
| Biclique c:9, p:5 | 1,969 | 12 | 0.63 | 15 | 0.13 | 50.64 | 1.24 |
| Biclique c:8, p:5 | 9,152 | 55 | 2.93 | 50 | 0.44 | 51.59 | 5.53 |
| Biclique c:8, p:4 | 16,345 | 88 | 4.68 | 75 | 0.65 | 53.61 | 8.55 |
| Biclique c:8, p:3 | 37,299 | 156 | 8.29 | 140 | 1.22 | 52.38 | 14.24 |

| | | | | | | | |
|---|---|---|---|---|---|---|---|
| Biclique c:7, p:4 | 54,119 | 241 | 12.83 | 200 | 1.73 | 54.63 | 20.68 |
| Biclique c:6, p:6 | 80,633 | 353 | 18.78 | 296 | 2.57 | 54.37 | 27.76 |
| Biclique c:7, p:3 | 97,031 | 342 | 18.23 | 320 | 2.78 | 51.57 | 26.83 |
| Biclique c:6, p:5 | 114,132 | 428 | 22.79 | 389 | 3.38 | 52.41 | 31.62 |
| Biclique c:6, p:4 | 166,065 | 520 | 27.69 | 517 | 4.49 | 50.13 | 35.55 |
| Biclique c:8, p:2 | 245,524 | 452 | 24.08 | 620 | 5.39 | 42.19 | 30.56 |
| Biclique c:6, p:3 | 257,527 | 636 | 33.87 | 734 | 6.38 | 46.46 | 39.09 |
| Biclique c:4, p:7 | 300,641 | 761 | 40.53 | 850 | 7.39 | 47.23 | 43.52 |
| Biclique c:5, p:5 | 355,552 | 814 | 43.34 | 977 | 8.49 | 45.43 | 44.25 |
| Biclique c:7, p:2 | 373,966 | 628 | 33.44 | 918 | 7.98 | 40.64 | 36.61 |
| Biclique c:4, p:5 | 464,224 | 904 | 48.13 | 1,200 | 10.43 | 42.96 | 45.32 |
| Biclique c:5, p:4 | 485,839 | 918 | 48.85 | 1,242 | 10.80 | 42.48 | 45.37 |
| Biclique c:4, p:4 | 600,488 | 990 | 52.70 | 1,470 | 12.78 | 40.24 | 45.57 |
| Biclique c:6, p:2 | 601,977 | 865 | 46.06 | 1,429 | 12.42 | 37.75 | 41.44 |
| Biclique c:5, p:3 | 683,556 | 1,032 | 54.92 | 1,630 | 14.16 | 38.76 | 45.39 |
| Biclique c:3, p:9 | 741,742 | 1,139 | 60.64 | 1,654 | 14.38 | 40.78 | 48.72 |
| Biclique c:4, p:3 | 795,442 | 1,079 | 57.45 | 1,839 | 15.98 | 36.99 | 44.96 |
| Biclique c:5, p:2 | 1,112,779 | 1,181 | 62.82 | 2,436 | 21.17 | 32.66 | 42.94 |
| Biclique c:4, p:2 | 1,254,992 | 1,227 | 65.31 | 2,694 | 23.42 | 31.30 | 42.29 |
| Biclique c:3, p:3 | 1,792,491 | 1,434 | 76.29 | 3,408 | 29.62 | 29.62 | 42.65 |
| Biclique c:2, p:9 | 2,014,916 | 1,539 | 81.87 | 3,415 | 29.68 | 31.07 | 45.04 |
| Biclique c:3, p:2 | 2,245,826 | 1,519 | 80.82 | 4,146 | 36.03 | 26.82 | 40.25 |
| Biclique c:2, p:2 | 4,740,224 | 1,776 | 94.48 | 6,998 | 60.83 | 20.25 | 33.34 |

***Supplementary Table 11.*** *KEGG enrichment for the E. coli input network (threshold 0.4) with p.adjust <=0.01, ordered by p.adjust.*

| Description | GeneRatio | BgRatio | p.adjust | fold.enrichment |
|---|---|---|---|---|
| Biosynthesis of secondary metabolites | 287/1076 | 339/1706 | 2.58E-20 | 1.34 |
| Biosynthesis of cofactors | 126/1076 | 142/1706 | 3.35E-11 | 1.41 |
| Biosynthesis of amino acids | 106/1076 | 117/1706 | 7.58E-11 | 1.44 |
| Microbial metabolism in diverse environments | 215/1076 | 267/1706 | 3.70E-10 | 1.28 |
| Carbon metabolism | 95/1076 | 110/1706 | 3.39E-07 | 1.37 |
| Alanine, aspartate and glutamate metabolism | 33/1076 | 33/1706 | 2.45E-06 | 1.59 |
| Pyrimidine metabolism | 48/1076 | 51/1706 | 2.45E-06 | 1.49 |
| Pyruvate metabolism | 53/1076 | 59/1706 | 2.56E-05 | 1.42 |
| Glycolysis / Gluconeogenesis | 43/1076 | 47/1706 | 6.99E-05 | 1.45 |
| Phenylalanine, tyrosine and tryptophan biosynthesis | 21/1076 | 21/1706 | 4.34E-04 | 1.59 |
| O-Antigen nucleotide sugar biosynthesis | 21/1076 | 21/1706 | 4.34E-04 | 1.59 |
| Folate biosynthesis | 25/1076 | 26/1706 | 6.32E-04 | 1.52 |
| Butanoate metabolism | 32/1076 | 35/1706 | 8.47E-04 | 1.45 |
| Arginine biosynthesis | 18/1076 | 18/1706 | 1.39E-03 | 1.59 |
| Fatty acid degradation | 15/1076 | 15/1706 | 5.24E-03 | 1.59 |
| Citrate cycle (TCA cycle) | 26/1076 | 29/1706 | 6.59E-03 | 1.42 |
| Propanoate metabolism | 32/1076 | 37/1706 | 6.59E-03 | 1.37 |
| beta-Alanine metabolism | 14/1076 | 14/1706 | 6.59E-03 | 1.59 |
| Biotin metabolism | 14/1076 | 14/1706 | 6.59E-03 | 1.59 |
| Pentose phosphate pathway | 28/1076 | 32/1706 | 8.08E-03 | 1.39 |
| Lysine biosynthesis | 13/1076 | 13/1706 | 9.06E-03 | 1.59 |
| One carbon pool by folate | 13/1076 | 13/1706 | 9.06E-03 | 1.59 |
| Purine metabolism | 61/1076 | 78/1706 | 9.06E-03 | 1.24 |

***Supplementary Table 12.*** *KEGG enrichment for the E. coli Biclique proteins c:5/p:2, p.adjust <= 0.01, n = 463 input proteins (threshold 0.4) with p.adjust <=0.01, ordered by p.adjust.*

| Description | GeneRatio | BgRatio | p.adjust | fold.enrichment |
|---|---|---|---|---|
| Ribosome | 36/299 | 78/1706 | 1.38E-07 | 2.63 |
| Pyrimidine metabolism | 22/299 | 51/1706 | 4.95E-04 | 2.46 |
| Citrate cycle (TCA cycle) | 15/299 | 29/1706 | 6.11E-04 | 2.95 |
| 2-Oxocarboxylic acid metabolism | 13/299 | 26/1706 | 2.60E-03 | 2.85 |
| Microbial metabolism in diverse environments | 67/299 | 267/1706 | 6.54E-03 | 1.43 |
| Pyruvate metabolism | 21/299 | 59/1706 | 7.07E-03 | 2.03 |
| Alanine, aspartate and glutamate metabolism | 14/299 | 33/1706 | 7.07E-03 | 2.42 |

***Supplementary Table 13.*** *KEGG enrichment for the E. coli Predicted proteins c:5/p:2, p.adjust <= 0.01, n = 420 input proteins (threshold 0.4) with p.adjust <=0.01, ordered by p.adjust.*

| Description | GeneRatio | BgRatio | p.adjust | fold.enrichment |
|---|---|---|---|---|
| Ribosome | 36/267 | 78/1706 | 4.35E-09 | 2.95 |
| Citrate cycle (TCA cycle) | 14/267 | 29/1706 | 1.21E-03 | 3.08 |
| Butanoate metabolism | 14/267 | 35/1706 | 9.84E-03 | 2.56 |

***Supplementary Table 14***. *Metabolites considered "currency metabolites" contained in the E. coli network.*

| KEGG ID | Chemical ID | Abbreviation |
|---|---|---|
| C00002 | CIDm00000238 | ATP |
| C00003 | CIDm00000925 | NAD+ |
| C00004 | CIDm00000928 | NADH |
| C00005 | CIDm00000930 | NADPH |
| C00006 | CIDm00000929 | NADP+ |
| C00008 | CIDm00000197 | ADP |
| C00010 | CIDm00000317 | CoA |
| C00013 | CIDm00001023 | Pyrophosphate |
| C00015 | CIDm00001158 | UDP |
| C00016 | CIDm00000703 | FAD |
| C00018 | CIDm00001051 | Pyridoxalphosphate |
| C00019 | CIDm00001079 | S-Adenosyl-L-methionine |
| C00020 | CIDm00000224 | AMP |
| C00021 | CIDm00000193 | S-Adenosylhomocysteine |
| C00035 | CIDm00000730 | GDP |
| C00044 | CIDm00000762 | GTP |
| C00055 | CIDm00000314 | CMP |
| C00061 | CIDm00000710 | FMN |
| C00063 | CIDm00000593 | CTP |
| C00075 | CIDm00001181 | UTP |
| C00105 | CIDm00001172 | UMP |
| C00112 | CIDm00000290 | CDP |
| C00113 | CIDm00001024 | PQQ |
| C00120 | CIDm00000253 | Biotin |
| C00144 | CIDm00000761 | GMP |
| C00175 | CIDm00002891 | Cobalt |
| C00194 | CIDm00002891 | Cobamidecoenzyme |
| C01352 | CIDm00000705 | FADH2 |

***Supplementary Table 15.*** *In relation to Supplementary Figure 3. Statistical results of biclique calculations for E. coli with different minimum number of compounds (c) and proteins (p) on each side of the biclique using a confidence threshold of 0.4 and allowing insertion of one edge to complete a biclique. The number of edges represents the number of predicted interactions. Datasets are sorted according to the number of edges. All values are the mean values of n=10 repetitions. Note that the underlying reference data was unbalanced with a mean number of positives n= 403 and a mean number of negatives n = 4,332.*

| Dataset | # Edges | # TP | TPR in % | # FP | FPR in % | PPV in % | F1 in % |
|---|---|---|---|---|---|---|---|
| Biclique c:8, p:4 | 32 | 27 | 6.71 | 1 | 0.02 | 97.38 | 12.53 |
| Biclique c:8, p:3 | 40 | 31 | 7.77 | 2 | 0.04 | 95.38 | 14.35 |
| Biclique c:6, p:6 | 44 | 28 | 6.83 | 2 | 0.05 | 92.97 | 12.70 |
| Biclique c:7, p:4 | 47 | 33 | 8.06 | 2 | 0.05 | 93.91 | 14.82 |
| Biclique c:8, p:2 | 52 | 38 | 9.50 | 2 | 0.05 | 94.57 | 17.26 |
| Biclique c:7, p:3 | 59 | 41 | 10.20 | 3 | 0.07 | 93.52 | 18.38 |
| Biclique c:6, p:5 | 63 | 35 | 8.56 | 4 | 0.09 | 90.24 | 15.59 |
| Biclique c:6, p:4 | 86 | 45 | 11.27 | 5 | 0.12 | 89.48 | 20.00 |
| Biclique c:7, p:2 | 100 | 54 | 13.30 | 6 | 0.13 | 90.56 | 23.18 |
| Biclique c:6, p:3 | 118 | 58 | 14.32 | 7 | 0.17 | 88.86 | 24.65 |
| Biclique c:6, p:2 | 187 | 75 | 18.66 | 10 | 0.24 | 87.80 | 30.76 |
| Biclique c:5, p:5 | 188 | 54 | 13.37 | 16 | 0.37 | 77.12 | 22.77 |
| Biclique c:5, p:4 | 245 | 67 | 16.70 | 19 | 0.44 | 77.93 | 27.48 |
| Biclique c:5, p:3 | 324 | 89 | 22.11 | 25 | 0.57 | 78.27 | 34.46 |
| Biclique c:5, p:2 | 457 | 110 | 27.19 | 32 | 0.74 | 77.48 | 40.18 |
| Biclique c:4, p:4 | 1,739 | 123 | 30.48 | 173 | 3.99 | 41.67 | 35.18 |
| Biclique c:4, p:3 | 2,138 | 145 | 35.84 | 208 | 4.80 | 41.04 | 38.25 |
| Biclique c:4, p:2 | 2,791 | 170 | 42.20 | 245 | 5.65 | 41.00 | 41.57 |
| Biclique c:3, p:3 | 20,056 | 253 | 62.74 | 1,668 | 38.49 | 13.27 | 21.88 |
| Biclique c:3, p:2 | 23,184 | 269 | 66.81 | 1,801 | 41.57 | 13.06 | 21.84 |
| Biclique c:2, p:2 | 92,731 | 386 | 95.78 | 3,780 | 87.25 | 9.27 | 16.90 |
